# Pou3f1 orchestrates a gene regulatory network controlling contralateral retinogeniculate projections

**DOI:** 10.1101/2022.06.28.497982

**Authors:** Michel Fries, Thomas W. Brown, Christine Jolicoeur, Camille Boudreau-Pinsonneault, Awais Javed, Pénélope Abram, Michel Cayouette

**Affiliations:** Cellular Neurobiology Research Unit, Institut de recherches cliniques de Montreal, Montreal, QC, H2W 1R7; Molecular Biology Program, Université de Montréal, Montreal, QC, H3C 3J7; Integrated Program in Neuroscience, McGill University, Montreal, QC, H3A 1A1

## Abstract

The balance of contralateral and ipsilateral retinogeniculate projections is critical for the establishment of binocular vision, but the transcriptional programs regulating this process remain ill-defined. In this study, we show that the Pou class homeobox protein POU3F1 is selectively expressed in developing mouse contralateral retinal ganglion cells (cRGCs). Inactivation of *Pou3f1* increases the number of ipsilateral RGCs produced, leading to abnormal ipsilateral to contralateral projection ratio at the optic chiasm, whereas expression of *Pou3f1* in retinal progenitors increases the production of cRGCs. Using Cut&Run and RNA sequencing in wildtype and *Pou3f1* mouse knockout retinas, we demonstrate that *Pou3f1* is necessary for the establishment of a cRGC gene regulatory network through activation and repression of several known contralateral and ipsilateral determinants, respectively. Finally, we report that POU3F1 is sufficient to induce production of RGC-like cells sending projections to the optic nerve, even in late-stage retinal progenitors not normally competent to generate RGCs. This work uncovers POU3F1 as a master regulator of the cRGC transcriptional program and opens new possibilities for optic nerve regenerative therapies.

## Introduction

The ability to perceive the world in 3D is critical to animal survival. This is known as binocular stereopsis and is mediated primarily by the ability to look at an object from two different parts of the retinas. The offset, and overlap, of the images produced from the left and right eye allows animals to accurately infer the depth and distance of an object in relation to themselves to make quick, and sometimes lifesaving judgements (Nityananda and Read, 2017; Patterson and Martin, 1992). Also critical to this process is the establishment of a proper ratio of retinal ganglion cells (RGCs) that cross the midline (contralateral RGCs; cRGC) and others that do not (ipsilateral RGCs; iRGCs) (Murcia-Belmonte and Erskine, 2019; Nityananda and Read, 2017; Patterson and Martin, 1992). Although the location of the eyes, either medially as in humans, or laterally as in rodents, generally correlates with the ratio of iRGCs/cRGCs, there are exceptions to this rule (Nityananda and Read, 2017), suggesting that molecular cues, rather than visual experience, are critical to establish the appropriate balance between ipsi- and contralateral projections.

The projection of RGCs is a well-coordinated process relying on a plethora of guidance cues and cell surface receptors that elicit an attractive or repulsive response in the growth cone, thereby guiding them towards the appropriate target. (Murcia-Belmonte and Erskine, 2019; Wang et al., 2016). While the various RGC axon guidance cues have been extensively studied (Kolodkin and Tessier-Lavigne, 2011; Wang *et al*., 2016), much less is known about the transcriptional programs enabling the appropriate expression of these effectors. One critical transcription factor is the zinc finger protein ZIC2, which has recently been established as a key determinant of the ipsilateral fate, in both the visual system and spinal cord (Escalante et al., 2013; Herrera et al., 2003; Matsuda et al., 2017). Additional studies have demonstrated that ZIC2 induces expression of the axon guidance receptor EPHB1 and the serotonin transporter SLC6A4 (Sert), which enable ipsilateral projection at the optic chiasm and refinement of eye-specific projections at retinorecipient targets, respectively (Garcia-Frigola et al., 2008; Garcia-Frigola and Herrera, 2010). On the other hand, the SOXC group of transcription factors (SOX4/11/12) and the LIM homeobox factor ISL2 have been identified as drivers of the contralateral fate (Kuwajima et al., 2017; Pak et al., 2004). ISL2 is exclusively expressed in cRGCs and *Isl2* knockout mice have more ipsilateral projecting ZIC2^+^ RGCs (Pak *et al*., 2004), suggesting that ISL2 ensures cRGC fate development by repressing iRGC determinants (Pak *et al*., 2004). SOXC are also expressed exclusively in cRGCs and knockdown of SoxC in the mouse retina results in reduced numbers of cRGCs (Kuwajima *et al*., 2017). It remains unknown, however, how exactly these determinants are activated in cRGCs during development. Interestingly, while expression of the bHLH transcription factor ATOH7 is rapidly turned off in differentiating cRGCs, it is maintained for some time in nascent iRGCs, suggesting that ATOH7 may be involved in controlling the cRGC/iRGC fate decision (Miesfeld et al., 2018; Wang *et al*., 2016). Knockout mice for *Atoh7* lack almost all RGCs, such that ATOH7 has long been thought to be the master regulator of RGC fate selection (Brown et al., 2001; Gao et al., 2014). But recent work has shown that inhibiting cell death in *Atoh7* deficient mice only results in a small decrease of total RGCs produced, indicating that ATOH7 is important for RGC survival, rather than fate specification (Brodie-Kommit et al., 2021). These results suggest that another, yet to be identified factor, controls the establishment of the cRGC transcriptional program in the developing retina.

Here we report that the POU class homeobox factor POU3F1 is a critical regulator of the iRGC/cRGC ratio determination. Among the developing RGC population, we show that POU3F1 is expressed only in cRGCs and inactivation of *Pou3f1* disrupts the proportion of axonal projections crossing the midline at the optic chiasm. Furthermore, ectopic expression of POU3F1 in retinal progenitors increases the generation of cRGCs at the expense of iRGCs. Using RNA-seq and Cut&Run sequencing, we show that POU3F1 binds to several known cRGC and iRGC determinants and upregulates or downregulates their expression, respectively, thereby ensuring proper iRGC/cRGC ratios. Finally, we show that expression of POU3F1 in late-stage progenitors that have normally lost the ability to generate RGCs is sufficient to induce production of RGC-like cells that send projections to the optic nerve. Expression of POU3F1 in also sufficient to rescue RGC production in *Atoh7* knockout mice. Together, our data identify POU3F1 as a master regulator of transcriptional programs underlying binocular vision in mammals.

## Results

### POU3F1 is expressed in differentiating contralateral retinal ganglion cells

To identify factors that might be involved in RGC subtype specification, we first analyzed previously published scRNA-seq data for transcripts that are enriched in mouse RGCs (Clark et al., 2019), particularly during the RGC production window between embryonic day (E) 12 and E16. We found that POU3F1 fits these criteria and is transiently expressed in developing RGCs at E14, which is the peak of RGC genesis, whereas it is downregulated at later stages (Fig 1A, Fig. S1). Additionally, Pou3f1 transcripts are strongly expressed in developing amacrine cells (Fig. 1A, Fig. S1). To examine POU3F1 protein expression, we next carried out immunostainings on mouse retinal sections from early developmental stages to adulthood. We found that 70-80% of POU3F1-positive cells are co-labelled with the RGC markers BRN3A (POU4F1) and BRN3B (POU4F2) at early embryonic stages (Fig. 1B-D). While POU3F1 expression is more prevalent in the BRN3B^+^ population at E13.5, it is reduced at later stages (Fig. 1C, D). Towards the end of the embryonic period and at early postnatal stages, when production of RGCs is largely complete, the proportion of POU3F1^+^ cells co-labelling with BRN3A or BRN3B decreases to less than 10% (Fig. 1C, D). These results indicate that POU3F1 is expressed in differentiating RGCs but is not generally maintained in mature RGCs (Miesfeld *et al*., 2018).

**Figure 1.**
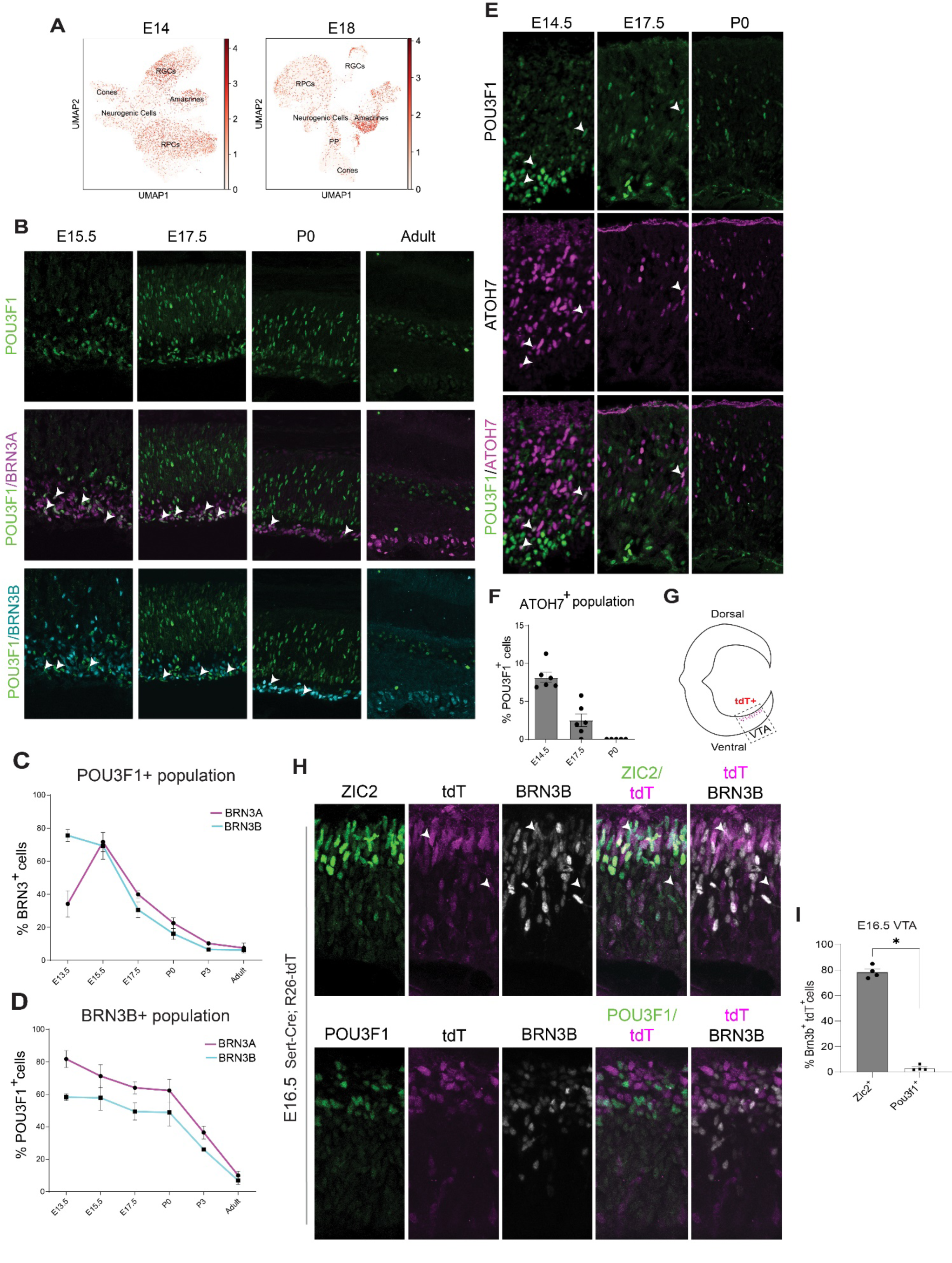
Expression of POU3F1 in the developing mouse retina. **(A)** scRNAseq UMAP plot of Pou3f1 expression at E14 and E18 in different retinal cell populations. **(B)** Co-immunostainings for BRN3A, BRN3B and POU3F1 in retinal sections at different stages of development in C57/Bl6 mice. White arrows point to cells showing co-localization of BRN3A/BRN3B and POU3F1. **(C)** Quantification of POU3F1^+^/BRN3A^+^ or POU3F1^+^/BRN3B^+^ at different stages of development (n = 5 animals per timepoint). **(D)** Quantification of BRN3A^+^ and BRN3B^+^ cell populations that are POU3F1^+^ (n = 5 animal per timepoint). **(E)** Co-immunostaining for POU3F1 and ATOH7. White arrows show co-labelled cells. **(F)** Quantifications of ATOH7^+^/POU3F1^+^ cells at E14.5, E17.5 and P0 (n = 4 animals per timepoint). **(G)** Schematic of Tdt expression in the ventro-temporal area (VTA) in Sert-Cre:R26-tdtomato mouse line. **(H)** Co-immunostaining for ZIC2 and BRN3B or POU3F1 and BRN3B in Sert-Cre:R26-tdtomato mouse line. White arrows point to co-labelled cells. **(I)** Quantification of the percentage of Brn3b+/Tdt+cells that also express ZIC2 or BRN3B in the VTA at E16.5.

To determine the identity of the remaining 20-30% of POU3F1^+^ cells that do not co-express BRN3A or BRN3B at early stages of retinal development, we asked whether POU3F1 might be expressed in neurogenic progenitors competent to generate RGCs. Most RGC-competent progenitors are defined by the expression of ATOH7 at embryonic stages (Brown *et al*., 2001; Gao *et al*., 2014). ATOH7 expression is largely restricted to progenitors throughout the retina from E13.5 to E17.5, and while it is downregulated in differentiating cRGCs, it is retained for a longer period in iRGCs (until about P0) (Wang *et al*., 2016). Immunostainings in the retina showed that about 8% of POU3F1^+^ cells are co-labelled with ATOH7 at E14.5 (Fig. 1E, F), and this proportion is further reduced at E17.5 and reaches 0% at P0 (Fig. 1F). Moreover, ATOH7^+^ cells mostly reside in the progenitor layer, whereas POU3F1^+^ cells mostly reside in the nascent ganglion cell layer (GCL) (Fig. 1E). Together, these results indicate that POU3F1 is expressed in a small fraction of RGC-competent progenitors and in the majority of differentiating RGCs.

Next, we asked whether POU3F1 might be expressed in a subtype of differentiating RGCs. As we found very few POU3F1^+^ cells co-expressing ATOH7, which is known to be retained for a prolonged period in iRGCs (Miesfeld *et al*., 2018; Wang *et al*., 2016), and POU3F1+ cells co-label with BRN3A, a cRGC marker (Quina et al., 2005; Wang *et al*., 2016). We suspected that POU3F1 may be specifically expressed in cRGCs. To investigate this, we relied on previously documented iRGC-specific markers, such as the ipsilateral determinant ZIC2 and the serotonin transporter SCL6A4 (Sert) to perform colocalization studies (Wang *et al*., 2016). We crossed a Sert*-*Cre driver mouse line (ET33 line) (Iwafuchi-Doi et al., 2012) with a Rosa26-tdTomato (R26-tdT) reporter line to generate a Sert*-*Cre; R26-tdT line in which iRGCs express the tdTomato (tdT) fluorescent reporter (Fig. 1G). As expected, a large proportion of the tdT^+^ cell population that co-expressed BRN3B in the ventrotemporal area (VTA), where iRGC are located, were also positive for ZIC2 at E16.5 (Fig. 1H, I) (78.4 ± 2.4%, n = 4). In contrast, we found very few POU3F1^+^ cells in the tdT^+^ /BRN3B^+^ population (Fig. 1H, I) (3.1 ± 1.0%, n = 4). These results indicate that POU3F1 is generally not expressed in developing iRGCs. Together with our observation that POU3F1 is found in cells expressing the cRGC marker BRN3A at E15-17 (Fig. 1C, D), these results show that POU3F1 labels cRGCs.

### POU3F1 promotes the cRGC fate

We next sought to determine whether POU3F1 controls the production of cRGCs, as suggested by our expression studies. To test this hypothesis, we first electroporated pCAG-IRES-GFP or pCAG-Pou3f1-IRES-GFP constructs in E14.5 retinal explants and six days later analysed the number of GFP+ cells that stained for the RGC markers BRN3A or BRN3B and the amacrine cell marker PAX6. We found about 10 times more GFP+ cells staining for the pan RGC marker BRN3B and for the cRGC marker BRN3A after POU3F1 expression, suggesting overproduction of cRGCs, but no difference in the number of PAX6+ cells in the INL, suggesting no change in amacrine cell production (Fig. 2A-C). Although most BRN3A/BRN3B+ cells produced after POU3F1 expression migrated to the ganglion cell layer, some ended up in other layers (Fig. 2A, B). Of note, we found a significant decrease in the number of electroporated cells residing in the retinal progenitor layer (RPL) in the *Pou3f1* condition, and more cells residing in the differentiated neuron layers (Fig. 2A, D), consistent with POU3F1 promoting differentiation. Moreover, we observed a significant decrease in the proportion of EdU^+^/GFP^+^ cells in *Pou3f1* electroporated explants that received a 2-hour pulse of EdU two days after electroporation (Fig. 2E, F), indicating that POU3F1 promotes cell cycle exit. These results show that expression of POU3F1 in retinal progenitors induces cell cycle exit and differentiation into cells expressing RGC markers.

**Figure 2.**
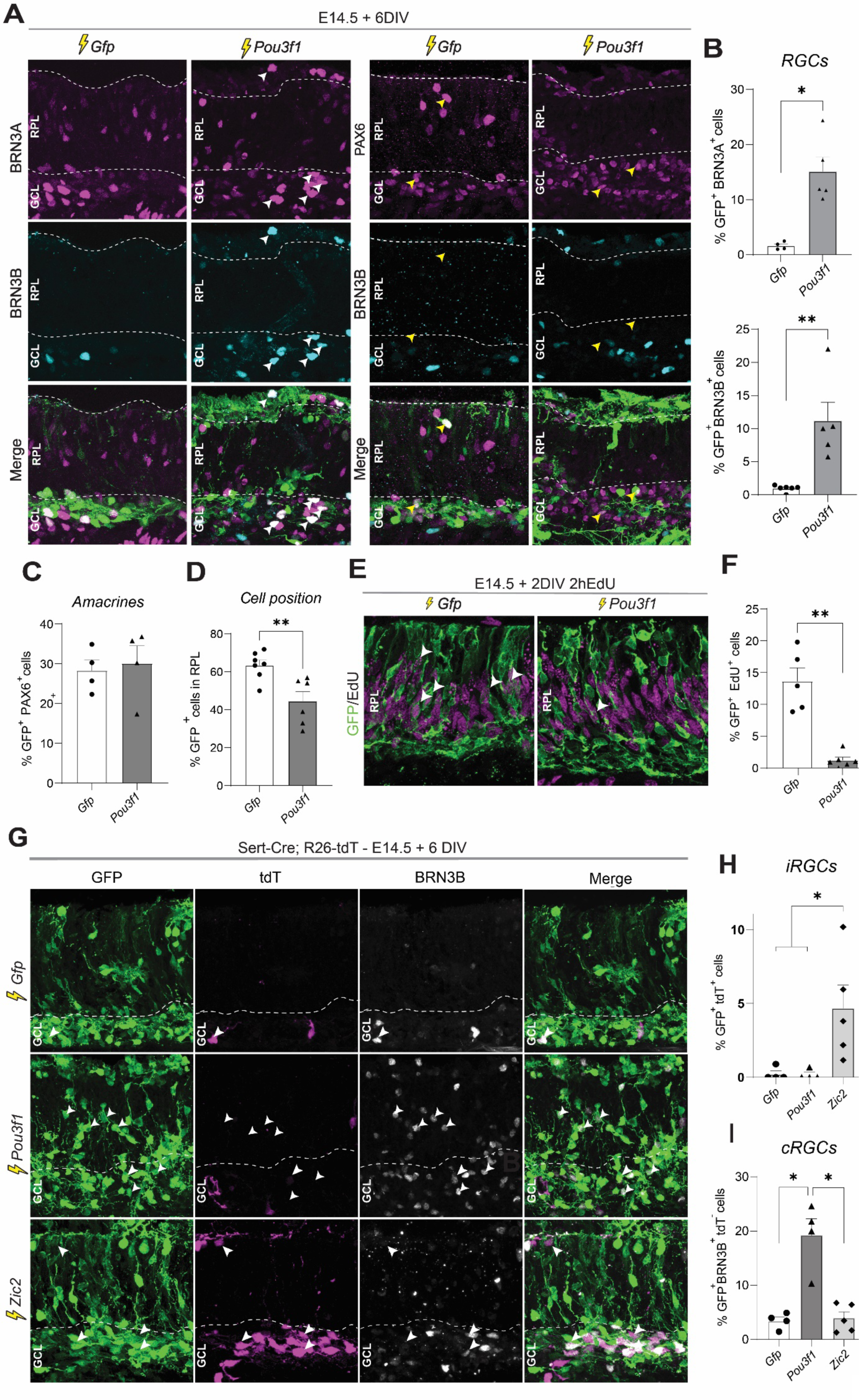
*Pou3f1* promotes the production of cRGCs. **(A)** Co-immunostainings on retinal cross sections from *Gfp* or *Pou3f1*-electroporated explants at E14.5 cultured for 6 days. Dotted line defines the retinal progenitor layer (RPL). White arrowheads point to triple positive cells for the listed markers, whereas yellow arrowheads point to PAX6+/BRN3B- cells. (**B)** Quantification of the proportion of electroporated cells (GFP+) that are BRN3A+ (top, n = 5) or BRN3B+ (bottom, n = 4) after GFP or POU3F1 overexpression. **(C)** Quantification of the proportion of electroporated cells (GFP+) expressing the amacrine cell marker Pax6. Only Pax6- positive/BRN3B-negative cells were counted to exclude RGCs (n = 4). **(D)** Quantification of the proportion of electroporated cells (GFP+) residing in the RPL. **(E)** Immunostaining for GFP and EdU on retinal explant sections 2 days after electroporation with GFP or POU3F1. EdU was added to the culture medium 2 hours before fixation. Arrowheads point out EdU^+^/GFP^+^ cells. **(F)** Proportion of GFP+ cells that are EdU+ in experiment shown in E (n = 5). **(G)** Co-immunostaining on retinal cross sections from GFP, Pou3f1 or Zic2-electroporated explants prepared from Sert-Cre;R26-tdT embryos at E14.5 and cultured for 6 days. Arrowheads point to triple positive cells for the listed markers (GFP^+^, TDT^+^, BRN3B^+^). **(H)** Quantification of *de novo* generated iRGC after electroporation with *Gfp* (n=4), *Pou3f1* (n=4) or *Zic2* (n=5) constructs. Multiple comparisons with one-way ANOVA and Tukey’s post-hoc test. **(I)** Quantification of the proportion of GFP^+^/BRN3B^+^/TDT^-^ cells after electroporation of *Gfp* (n=4), *Pou3f1* (n=4) or *Zic2* (n=5) constructs. Multiple comparisons with one-way ANOVA and Tukey’s post-hoc test.

To more precisely identify POU3F1-induced cells, we next electroporated retinas from Sert-Cre; R26-tdt embryos at E14.5 with either pCAG-IRES-GFP alone (*GFP*) as control, pCAG-Pou3f1-IRES-GFP (*Pou3f1*), or pCAG-Zic2-IRES-GFP (*Zic2*), which was previously reported to promote production of iRGC, and cultured explants for six days (Herrera *et al*., 2003). In this assay, we defined GFP^+^/TDT^+^ double positive cells as *de novo-*generated iRGCs, and GFP^+^/BRN3B^+^/TDT^-^ as de novo-generated cRGCs. In the *GFP* condition, we observed only a few iRGCs produced (Fig. 2A, B), as expected given that only a small population of progenitors give rise to iRGCs and they are restricted to the ventro-temporal area (VTA). After *Zic2* electroporation, we observed a significant increase in iRGC production (Fig. 2G, H), as previously reported (Wang *et al*., 2016). Interestingly, after electroporation of *Pou3f1,* we observed no change in iRGCs produced but found a 5-fold increase in GFP+/TDT- cells that stain for BRN3B (Fig. 2G-I). Together, these results suggest that POU3F1 promotes acquisition of the cRGC fate.

### Pou3f1 is necessary for cRGC production

Based on the above results, we predicted that POU3F1 may be required for newborn RGCs to acquire a contralateral identity. To elucidate the role of POU3F1 in RGC projections, we studied *Pou3f1* knockout mice. Inactivation of *Pou3f1* was previously reported to disrupt Schwann cell development and myelination in the peripheral nervous system (Bermingham et al., 1996; Jaegle et al., 2003). More recently, *Pou3f1* was found to induce neuronal gene expression in epiblast stem cells, but the role of *Pou3f1* in the central nervous system in vivo remains unknown (Zhu et al., 2014). To study *Pou3f1* function in the retina, we used a previously published *Pou3f1* KO mouse line (*Pou3f1*^lacZ/lacZ^), which carries a *lacZ* reporter knocked-in the POU-domain of *Pou3f1*, resulting in the production of a truncated non-functional POU3F1 protein (Jaegle *et al*., 2003). Consistently, we detected a truncated POU3F1 protein in the retina of *Pou3f1* KO mice (Fig. S2). Given that 98% of *Pou3f1* KO are perinatally lethal (Jaegle *et al*., 2003), we crossed *Pou3f1*^lacZ/+^ (HET) mice to generate *Pou3f1* KO and analyzed tissues at embryonic stages. We first asked whether generic RGC production is altered in absence of POU3F1. To do this, we stained *Pou3f1* KO retinas for the RGC markers BRN3A and ATOH7 at E14.5 (Fig. 3A). We found that *Pou3f1* KO retinas have similar numbers of ATOH7^+^ or BRN3A^+^ cells as the *Pou3f1* HET (Fig. 3B), suggesting that loss of POU3F1 does not affect overall generation of RGCs. To explore if loss of POU3F1 more specifically affects the cRGC/iRGC ratio, we stained E17.5 retinas for ATOH7, which is maintained in iRGCs at this stage, and for the iRGC-specific marker ZIC2. As predicted, we found that the populations of ATOH7^+^ and ZIC2^+^ cells are increased in *Pou3f1* KO compared to *Pou3f1* HET retinas, both in the central retina and peripheral VTA (Fig. 3C-F).

**Figure 3.**
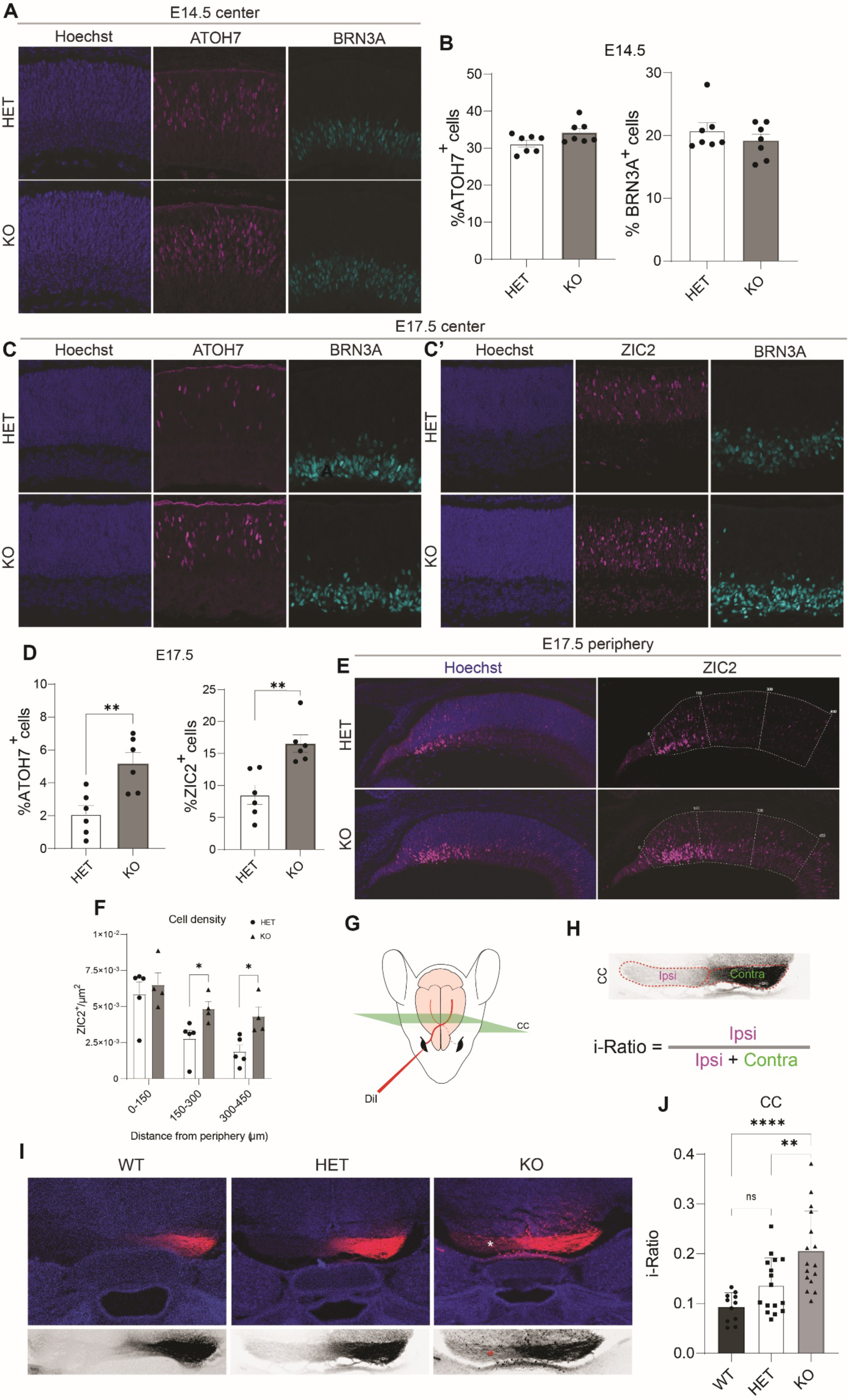
Loss of *Pou3f1* causes ectopic projection ratios at the optic chiasm. **(A)** Representative images from central retina at E14.5 from *Pou3f1*^+/-^ (HET) and *Pou3f1*^-/-^ (KO) mice stained for ATOH7, BRN3A and Hoechst nuclear dye. **(B)** Quantification of the proportion of ATOH7+ and BRN3A+ cells in central retina of HET vs KO at E14.5 (n=7) **(C)** Representative images from the central retina of HET and KO mice at E17.5 stained for ATOH7 and BRN3A (left) and ZIC2 and BRN3A (right). Cell nuclei were stained with Hoechst. **(D)** Quantification of the proportion of ATOH7+ and ZIC2+ cells in central retina of HET (n=6) vs KO (n=6) mice at E17.5. **(E)** Representative images of the peripheral retina of HET and KO mice at E17.5 stained for Zic2 (magenta). Retinal periphery was subdivided into three areas (dotted fields) that have the same width (150 µm). **(F)** Quantification of the number of Zic2+ cells/µm2 in each area highlighted in (F) in HET (n = 4) and KO (n = 4). Analyzed surface areas were similar in size among all samples and did not display any significant deviation between repeats. **(G)** Schematic representation of the experimental procedure. DiI was injected in the right eye of E17.5 embryos and heads were sectioned coronally 10 days later. The green square represents the region of interest located immediately after the optic chiasm (OC) midline crossing. This area was defined as the caudal chiasm (CC). **(H)** Representative image of the CC 10 days after DiI injection in the right eye. Fluorescence intensities were measured in the ipsi- and contralateral tracts (red dotted lines) and i-Ratio was calculated using the listed formula. **(I)** Representative images of the CC from WT, HET and KO mice stained for DiI (red) and Hoechst. The asterisk in the ipsilateral tract of the KO highlights the presence of a stronger DiI intensity compared to the HET and WT sample. Bottom panels show black and white images of the zoomed-in CC region shown in the respective panels above. **(J)** Graph of i-Ratio calculations in WT, HET and KO mice. (WT: n = 11; HET: n = 16; KO: n = 16; **p< 0.01, **** p<0.0001, multiple comparisons with one-way ANOVA and Tukey’s post-hoc test).

Based on the above results, we predicted that loss of POU3F1 would alter the ratio of contralateral and ipsilateral RGC projections. To explore this possibility, we retrogradely labeled RGC axons using DiI at E17.5, as previously reported (Garcia-Frigola *et al*., 2008; Garcia-Frigola and Herrera, 2010; Herrera *et al*., 2003). We then sectioned the heads transversely and quantified the proportion of iRGC to cRGC projections (defined here as i-Ratio) by measuring fluorescence intensity at the caudal chiasm (CC), just after midline crossing (Fig. 3G, H), as previously described (Peng et al., 2018). Although we observed a trend for an increase in i-Ratio in HET compared to WT, the difference was not significant (Fig. 3I, J). In Pou3f1 KO mice, however, the i-Ratio was significantly increased, with more labeled fibers in the ipsilateral tract (Fig. 3I, J). Taken together, these results show that Pou3f1 is required to establish the proper iRGCs/cRGCs projection ratio.

### POU3F1 regulates the cRGC transcriptional program

To unravel the genetic programs by which POU3F1 promotes the cRGC fate, we carried out transcriptional profiling experiments. We first electroporated retinas at P1 with *Gfp* or *Pou3f1 in vivo*, sacrificed the animals 48 hours or 14 days later, and sorted GFP^+^ cells for RNA extraction and sequencing. We decided to express *Pou3f1* at P1 rather than E14.5 because this is a time when RGCs are not normally produced and *Pou3f1* is no longer expressed, making it easier to identify transcripts regulated by POU3F1.

We started by addressing the rapid transcriptional response induced by POU3F1 by analyzing the transcriptome 48 hours after expression. We found 940 up-regulated (*Log2FC > 0.25, P-value < 0.05*) and 1250 down-regulated (*Log2FC < -0.25, P-value < 0.05*) genes (Fig. 4A, Table S1). By Gene ontology (GO) enrichment analysis, upregulated genes were associated with nervous system development, cell-projection organization, head and brain development (Fig. 4B), whereas downregulated genes were associated with cell differentiation, regulation of cell differentiation, neuron and eye development (Fig. 4C), among others. The most significantly up-regulated genes were *Eomes* and *Alcam* (Fig. 4A), which were shown to play essential roles in RGC axonogenesis (Mao et al., 2008; Thelen et al., 2012). Other notable upregulated transcripts included *Cdh6, 7, 8, 10, 18* and *24* (Fig. 4A), which code for a large family of cell adhesion proteins involved in dendrite growth of ON-OFF direction-selective RGCs (Duan et al., 2018). Genes coding for presynaptic proteins normally expressed in RGCs such as *Nrxn1*, *Slitrk1/2* and *Lrfn5* were also upregulated (Wang *et al*., 2016). Among the list of most down-regulated transcripts, we found genes involved in ipsilateral axon guidance, such as *EphB1* and *Fzd8* (Fig. 4A) (Garcia-Frigola *et al*., 2008; Lee et al., 2008; Wang *et al*., 2016), and in the regulation of proliferation such as *Notch1* and *Hes5*, which are also repressed by the SOXC transcription factors (Fig. 4A) (Kuwajima *et al*., 2017). Interestingly, we also found that *Atoh7* is down regulated 48 hours after *Pou3f1* expression (Fig. 4A).

**Figure 4.**
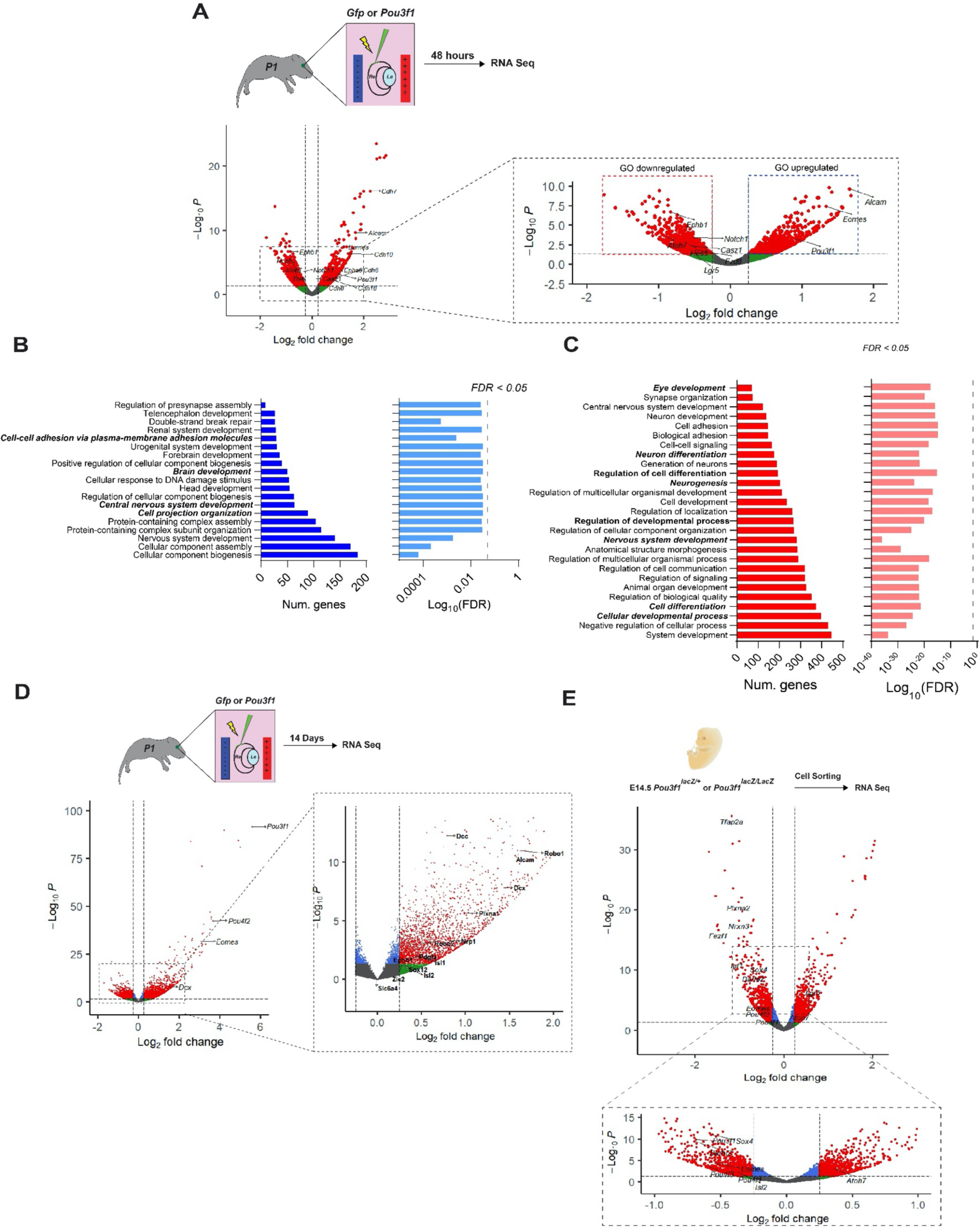
*Pou3f1* induces a cRGC transcriptional program. **(A)** Volcano plot of differentially expressed genes 48 hours after Pou3f1 expression in P1 retina. Red dots represent significantly up- or downregulated hits (Log2.FC > |0.25|; *p-value < 0.05*). Both up- and downregulated portions were used to perform GO term analysis (Log2.FC: Log2 fold change). **(B, C)** GO analysis of upregulated (B, blue) and ydownregulated (C, red) genes. Left bar plots show the number genes assigned to a biological process. Right bar plots represent the significance of enrichment with a false discovery rate (FDR) set at 0.05 (dotted black line). **(D)** Volcano plot of differentially expressed genes 14 days after expression of Pou3f1 in P1 retina. **(E)** Volcano plot of differentially expressed genes in FACS-sorted ß-Gal+ cells from Pou3f1^LacZ/+^ compared to Pou3f1^LacZ/LacZ^ E14.5 retinas. Red dots represent significantly up- or downregulated transcripts (Log2.FC > |0.25|; *p-value < 0.05*).

Next, we compared the transcriptome of *Pou3f1*-expressing cells with that of *Gfp* control cells 14 days after electroporation. We identified 2415 upregulated genes in the *Pou3f1* condition (*Log2FC > 0.25, P-value < 0.05*), which contained key effectors of RGC development and cRGC fate specification, such as BRN3B (*Pou4f2*) and DCX, consistent with our immunostaining results, as well as *Alcam* and *Eomes* (Fig. 4D, Table S2). Interestingly, sustained expression of *Pou3f1* until P14 led to the induction of the cRGC determinants Sox4 and 11 (Fig. 4D, dotted box), but not *Isl2* (*Log2FC = 0.32; P-value = 0.1*) nor Sox12 (*Log2FC = 0.247, P-value = 0.14*) (Kuwajima *et al*., 2017; Lee *et al*., 2008; Pak *et al*., 2004). We also observed increased expression of the guidance receptors *Nrp1* and *PlexinA1*, which are necessary for midline crossing at the optic chiasm (Fig. 4D, dotted box) (Andermatt et al., 2014; Erskine et al., 2011; Kuwajima et al., 2012). Furthermore, we found induction of the pan-RGC guidance receptors *Robo1*, *Robo2* and *DCC* (Deiner et al., 1997; Erskine et al., 2000; Ringstedt et al., 2000), and *Pdgfα*, which codes for platelet-derived growth factor secreted by endogenous RGC to allow expansion of astrocytes during retinogenesis (Fig. 4D, dotted box) (Fruttiger et al., 1996). Finally, we observed no changes in the levels of the ipsilateral markers *Zic2*, *EphB1* and *Sert* (SLC6A4), as expected given our findings that POU3F1 is required for the cRGC fate.

To identify the transcriptional programs normally controlled by POU3F1 in the developing retina, we next compared the transcriptome of *Pou3f1* lineage cells expressing *Pou3f1* or not. To do this, we took advantage of the *lacZ* insertion at the *Pou3f1* locus in Pou3f1^lacZ/lacZ^ mice. We sorted ß-Gal^+^ cells from *Pou3f1* KO and *Pou3f1* HET animals at E14.5 and compared their transcriptome by RNA-seq. We found that loss of POU3F1 leads to a significant down-regulation (P < 0.05; Log2FC < -0.25) of several factors involved in cRGC development such as *Igf1*, *Plxna2*, *Pou4f1*/*Brn3a*, *Isl2*, *Sox4*, *Fezf1* and *Nrxn3* (Fig. 4E, Table S3) (Kuwajima *et al*., 2017; Wang *et al*., 2016).

To more extensively analyze genes associated with the cRGCs fate, we cross-referenced our *Pou3f1* KO RNA-seq dataset with a previously published microarray dataset of cRGCs carried out at E16.5 (Wang *et al*., 2016). If POU3F1 is indeed a cRGC determinant, we expected that genes previously identified as enriched in cRGCs would be downregulated in *Pou3f1* KO cells. Consistent with this idea, we found that 14 out of 38 cRGC genes (36.8%) such as *Sema3e*, *Igf1*, *Rgs4* and *Isl2* are significantly downregulated in *Pou3f1* KO and only three are upregulated (Fig. S3A). Next, we compared transcriptional changes in *Pou3f1* KO cells to those observed after *Zic2* overexpression at E14.5, which promotes iRGC and reduces cRGC (Escalante *et al*., 2013). We predicted that the list of downregulated transcripts observed after inactivation of *Pou3f1* or *Zic2* overexpression would overlap. Consistently, out of the 643 down- regulated genes in the *Zic2* overexpression dataset, 21% were also significantly reduced in the *Pou3f1* KO (Fig. S3B) (Escalante *et al*., 2013). Notably, the list of genes included *Pou3f1* itself, as well as *Sox4*, *Apc2*, *Barhl2*, *Rgs4*, *Tfap2a* and *Chrnb3*, which was the strongest downregulated gene in the *Pou3f1* KO and its expression was recently reported in RGCs and retinorecipient areas of the brain (Drayson and Triplett, 2019).

To filter out the POU3F1 positively and negatively regulated genes, we cross-referenced this *Pou3f1* KO RNA-Seq dataset with that of *Pou3f1* overexpression at 48h (Fig. S3C- F). Among the positively regulated genes (i.e. upregulated in *Pou3f1* overexpression and down-regulated in *Pou3f1* KO), we found *Eomes*, *Nrxn1*, *Slitrk1/2* and *Tet2* (Fig. S3D, G). We obtained fewer candidates for negatively regulated genes (i.e., downregulated in *Pou3f1* overexpression and unregulated in *Pou3f1* KO) but, interestingly, *Atoh7* was in this group (Fig. S3F, H), as well as *Hes5* and *Notch1* (Fig. S3H), which are repressed by SOXC to promote the cRGC fate (Kuwajima *et al*., 2017).

The above results suggest that POU3F1 activates the cRGC transcriptional program, while repressing iRGC determinants like *Atoh7* and *Zic2*. Consistently, when we expressed *Pou3f1*-IRES-GFP in E14.5 retinas and stained for ATOH7 and ZIC2 48 hours later, we observed a drastic reduction in the number of ATOH7^+^/GFP^+^ and ZIC2^+^/GFP^+^ cells in the VTA compared to GFP expression alone (Fig. 5A-C).

**Figure 5.**
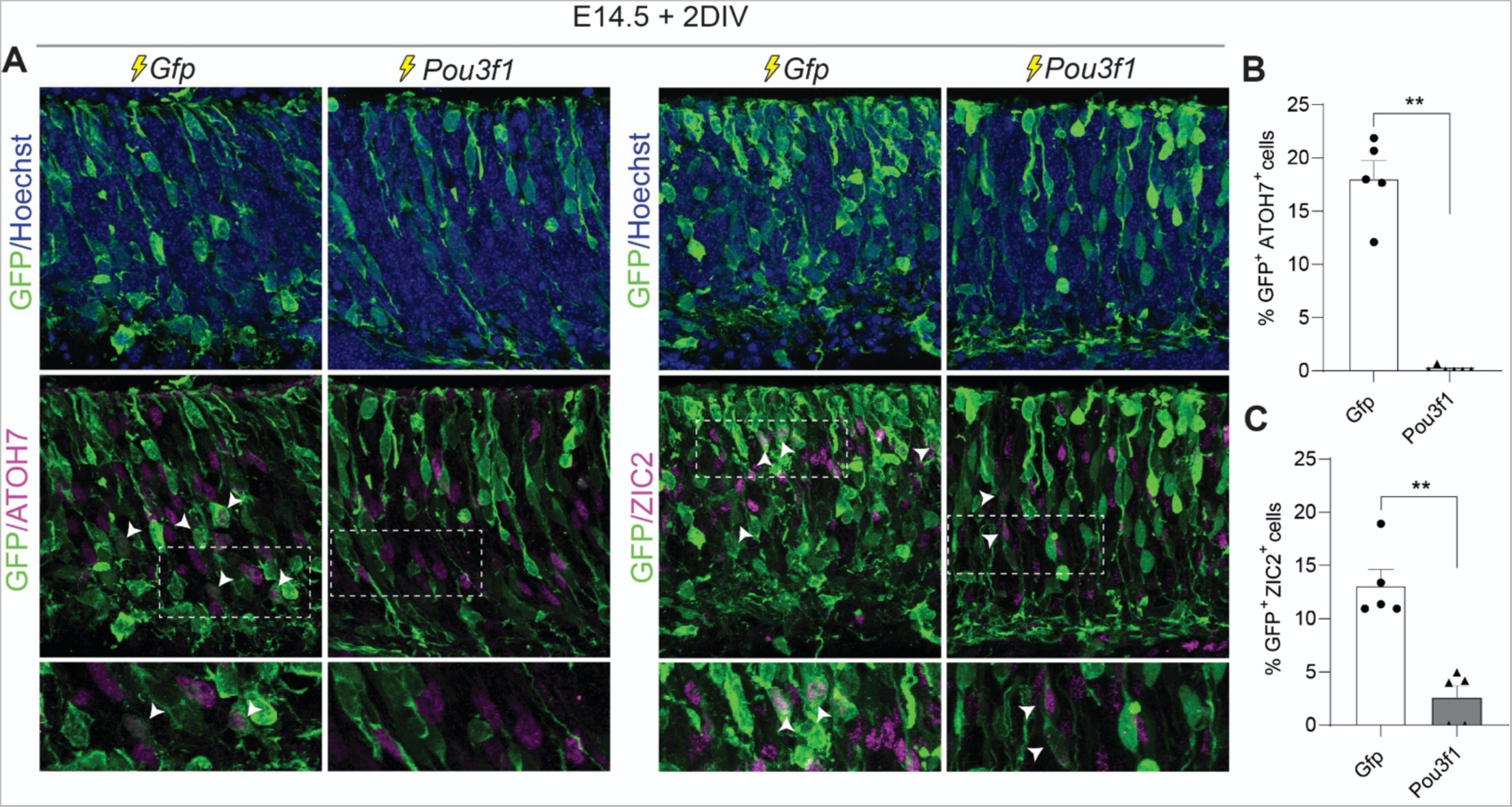
*Pou3f1* expression reduces ATOH7- and ZIC2-expressing cells. **(A)** Representative images of retinal explant cross sections two days after electroporation of *GFP* or *Pou3f1* constructs at E14.5. Sections were stained for either ATOH7 (left panels) or Zic2 (right panels). White arrowheads highlight ATOH7^+^/GFP^+^ or ZIC2^+^/GFP^+^ cells. (**B, C)** Quantification of ATOH7^+^/GFP^+^ cells (B, n=5) or ZIC2^+^/GFP^+^ (C, n = 5) from the total GFP^+^ population in *Gfp* or *Pou3f1* electroporated explants.

### POU3F1 binds cis regulatory elements of genes involved in RGC fate determination

To identify POU3F1 bound regions across the genome, we carried out CUT&RUN sequencing on E14.5 wildtype retinas and cross-referenced the list of bound genes to that of differentially expressed genes in the *Pou3f1* KO cells identified by RNA-seq (Fig. 6A). We found a total of 847 genes that were both bound by POU3F1 and differentially expressed in *Pou3f1* KO cells (Fig. 6A, Table S3). To obtain a general idea of the function of POU3F1 bound genes, the CUT&RUN peaks were analyzed with Genomic Regions Enrichment of Annotations Tool (GREAT) and classified using the GO Mouse Phenotype function (McLean et al., 2010). We observed that the strongest hit was *perinatal lethality*, followed by *abnormal nervous system*, *optic-tract morphologies*, *optic nerve hypoplasia* and *abnormal GCL morphology* (Fig. 6B), consistent with our findings that POU3F1 regulates cRGC production. Binding motif analysis identified the canonical POU-binding domain ‘TGCAAAT’ as the second ranked motif with the lowest amount of background (Fig. 6C). Importantly, the overall genome-wide distribution of POU3F1 observed by CUT&RUN is consistent with that observed in a previously published ChIP-Seq study in epiblast stem cells (Matsuda *et al*., 2017), validating the assay (Fig. 6D). Next, we cross-referenced the gene subsets related to *abnormal GCL morphology* and *abnormal optic nerve morphology* in the CUT&RUN GO Mouse Phenotype analysis against the *Pou3f1* KO RNA-Seq data to determine which POU3F1 bound genes are most significantly downregulated (Log2FC < -0.25; P-value < 0.05;) after *Pou3f1* inactivation (Fig. 6E, F). We observed that the expression of one of the SoxC genes, Sox4, is reduced in *Pou3f1* KOs and contains POU3F1 binding peaks adjacent to the promoter (Fig. 6G).

**Figure 6.**
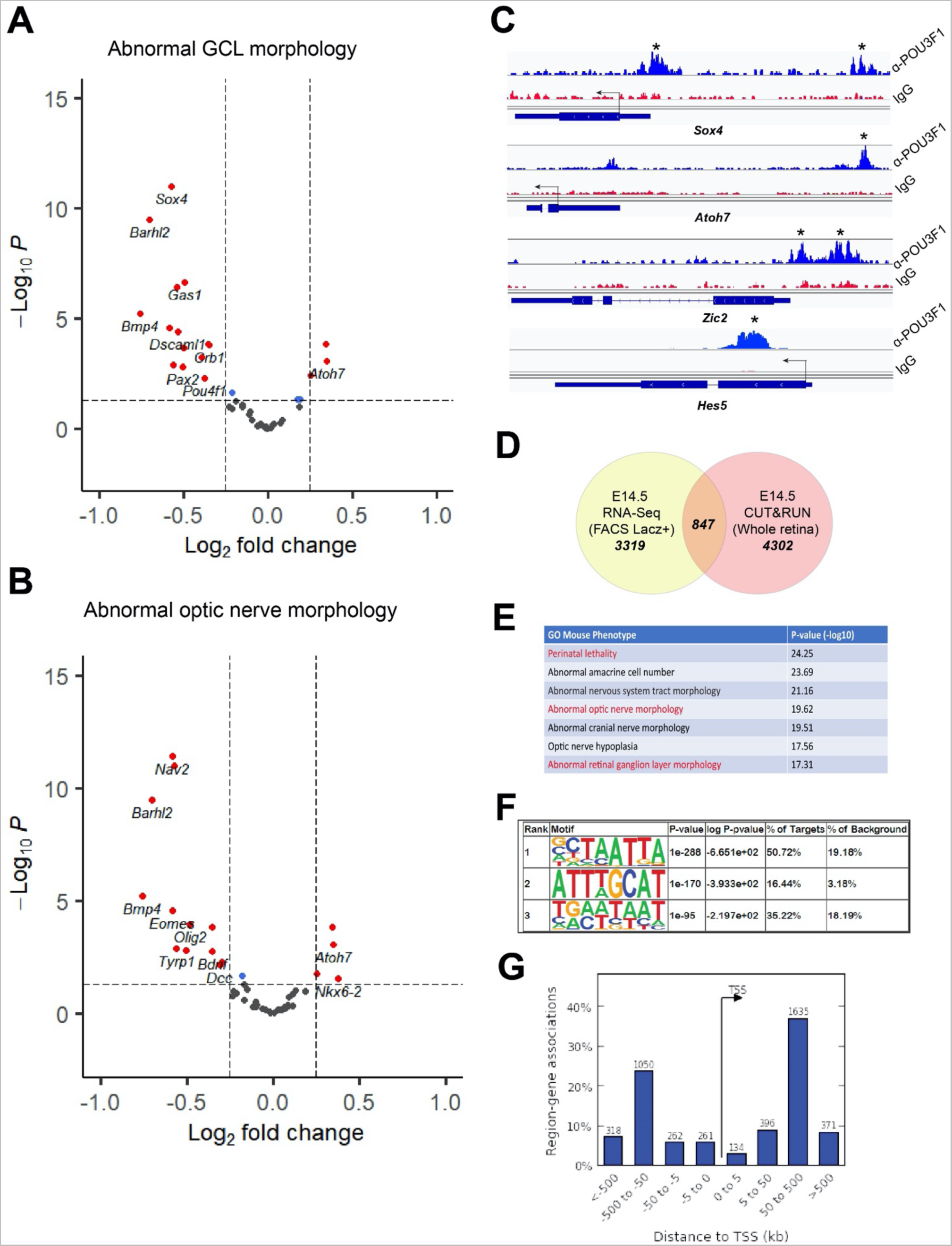
POU3F1 genome-wide binding profile. **(A, B)** Volcano plots of DEGs bound by POU3F1 listed in the GO terms: *Abnormal GCL morphology* (A) and *Abnormal optic nerve morphology* (B). Most of the genes associated with these terms are downregulated. **(C)** Binding profile of POU3F1 at *Sox4* and *Atoh7* gene regions in E14.5 retinas. Asterisks highlight significant binding peaks with a FDR < 0.05 (α-Pou3f1: Pou3f1 specific antibody; IgG: IgG-isotype control antibody). **(D)** Venn Diagram showing the number of differentially expressed genes (DEGs) in the E14.5 *Pou3f1* KO RNA seq (see Fig. 4E) and the number of bound genes in the E14.5 CUT&RUN-Seq. A total of 847 genes are bound by POU3F1 and differentially expressed in *Pou3f1* KO. **(E)** Table listing the top GO terms associated to the bound genes found in the POU3F1 CUT&RUN. **(F)** Binding motifs analysis showing the canonical POU-binding sequence as the second rank hit with the lower percentage of background. **(G)** The genome-wide binding distribution of POU3F1. Preferential binding is observed distally (500 to 50kB or -500 to -50kB) to the transcription start site (TSS), as previously reported (Matsuda *et al*., 2017).

Furthermore, other bound genes such *Barhl2* and *Nav2* are downregulated in *Pou3f1* KO (Fig. 6E, F). Interestingly, *Barhl2* was found to regulate contra- versus ipsilateral fate acquisition in developing dl1 neurons (Ding et al., 2012), and *Nav2* is necessary for cranial nerve development and neurite outgrowth (McNeill et al., 2010; Panza et al., 2015). Finally, we found that POU3F1 binds to a region upstream of the *Atoh7* and *Zic2* promoters (Fig. 6G), consistent with our observation that both *Atoh7* and *Zic2* are downregulated after *Pou3f1* overexpression (Fig. 4A, D), and that *Atoh7* is upregulated in *Pou3f1* KO (Fig. 4E). Together with our RNAseq results, this data indicates that POU3F1 binds to cis regulatory elements of several contralateral and ipsilateral fate determinants, upregulating or downregulating their expression, respectively. This leads to the establishment of a gene regulatory network enabling generation of the proper cRGC/iRGC ratio.

### POU3F1 is sufficient to promote RGC-like cell production from late-stage progenitors in an ATOH7-independent manner

The identification of novel RGC-promoting factors opens exciting possibilities to develop cell therapies for degenerative diseases of the optic nerve, particularly if these factors are sufficient to trigger the RGC fate. To study this question, we ectopically expressed POU3F1 in late retinal progenitors, which have normally lost the competence to generate RGCs. We first electroporated either *Gfp* or *Pou3f1* expression constructs in 1-day old postnatal retinas (P1) and cultured them as explants for 9 days (Fig. S4A). As endogenous RGCs have their axons severed during postnatal retinal explant preparation, they undergo rapid cell death in culture (Manabe et al., 2002), rendering the identification of de novo RGCs easy to monitor by marker expression and appearance of axonal outgrowth from the explants. In *Gfp*-electroporated explants, we found no axonal outgrowth during the entire culture period, as expected (Fig. S4B, top panels). In contrast, in explants electroporated with *Pou3f1*, we found clear axonal outgrowth as early as 4 days in vitro (DIV) that became denser and more complex by 9 DIV (Fig S4B, lower panels), suggesting de novo RGC production from late-stage progenitors. To substantiate this conclusion, we sectioned the explants and stained for RGC markers. We observed a significant increase in GFP^+^/BRN3B^+^ cells in *Pou3f1* electroporated explants (Fig. S4 C, D). Interestingly, however, we found that these cells do not express other markers of RGCs such as BRN3A or ISL1 (both 0%; n = 3). We therefore termed those ectopic BRN3B^+^ cells as RGC-like (RGC-L) cells.

To address if RGC-L cells arise *de novo* from progenitors or whether they might be surviving endogenous RGCs, we carried out a 2h EdU pulse after electroporating P0 retinal explants (EdU 0DIV) and continued the cultures for 7 days (Fig. S5A). No difference was observed in the GFP^+^/EdU^+^ populations between the control and *Pou3f1* condition (Fig S5B-D), indicating that, unlike what we observed at E14.5, POU3F1 does not trigger premature cell cycle exit when expressed in progenitors at P0. However, we noticed that more than 15% of the *Pou3f1* electroporated cell population was EdU^+^/BRN3B^+^ cells, whereas none of the control GFP transfected cells were EdU^+^/BRN3B^+^ (Fig. S5B-E). Moreover, when performing the EdU pulse 1 day after electroporation (EdU 1DIV), we observed an important reduction of GFP^+^/EdU^+^ cells in the *Pou3f1* condition (Fig. S5F). Taken together, these results show that POU3F1 induces *de novo* RGC-L cell production from postnatal progenitors and suggest that the RGC-L fate is acquired rapidly after cell cycle exit.

To test if POU3F1 could similarly promote RGC-L cell production *in vivo,* we electroporated *Gfp* or *Pou3f1* expressing constructs in P1 animals and collected the eyes 21 days later (Fig. 7A). We first evaluated the abundance of GFP^+^/BRN3B^+^ and GFP^+^/DCX^+^, two pan-RGC markers, in both control and *Pou3f1* conditions three weeks after electroporation (Fig. 7A). We found that control retinas have no GFP^+^/BRN3B^+^ and very few GFP^+^/DCX^+^ cells, which is also expressed in mature horizontal cells (Wakabayashi et al., 2008). In contrast, we found many GFP^+^/BRN3B^+^ and GFP^+^/DCX^+^ cells in *Pou3f1* electroporated retinas (Fig. 7B-D), consistent with our findings in explants. These ectopic RGC-L cells were generated at the expense of retinal cell types normally produced at postnatal stages, as the proportions of photoreceptors (GFP^+^ cells in ONL), bipolars (GFP^+^/CHX10^+^) and Müller glia (GFP^+^/LHX2^+^) cells were reduced after expression of *Pou3f1* compared to control (Fig. S6).

**Figure 7:**
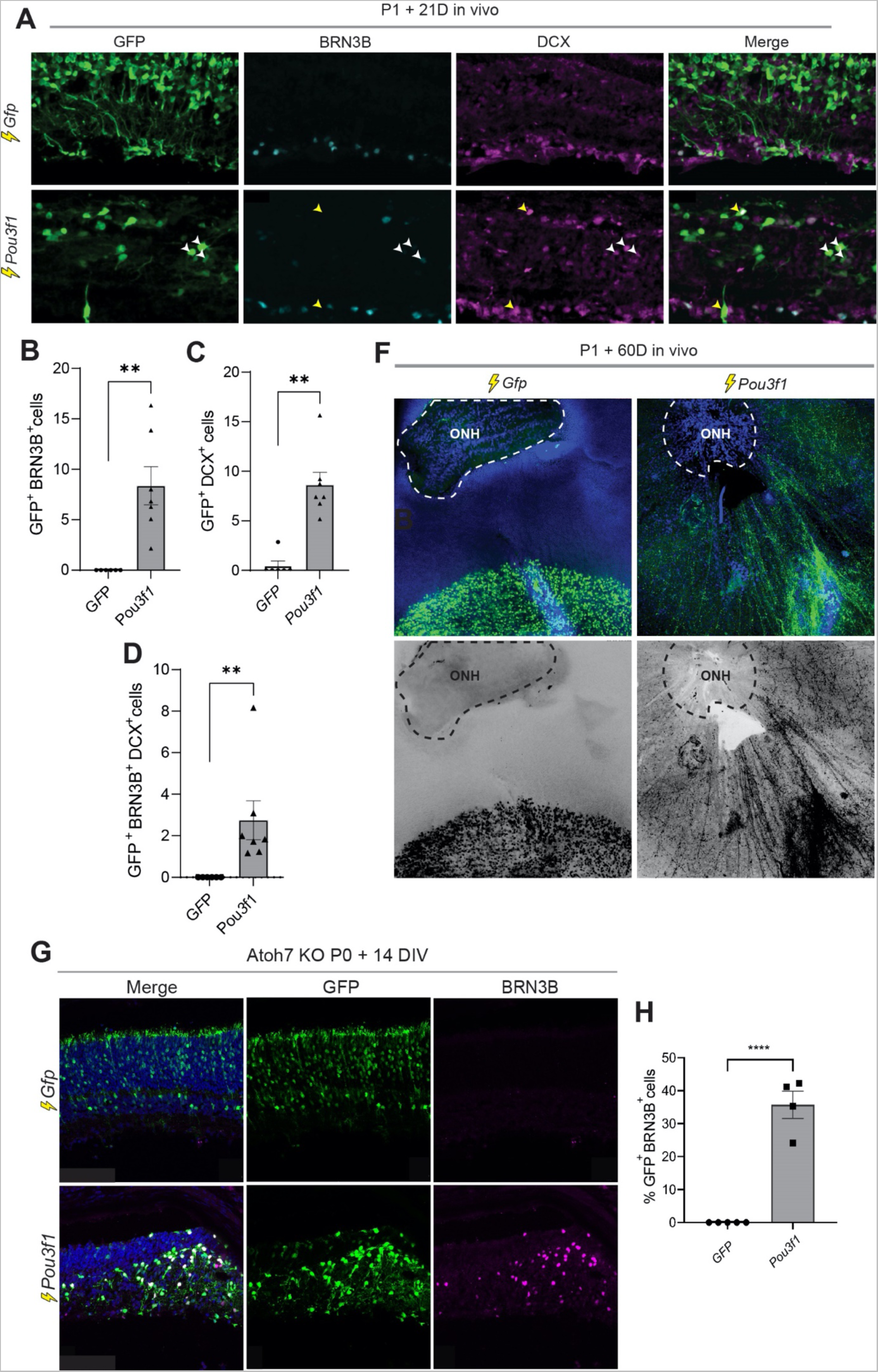
Misexpression of *Pou3f1* in the postnatal retina induces the production of RGC-like cells in an ATOH7-independent manner. **(A)** Immunostaining for BRN3B and Doublecortin (DCX) on retinal cross-sections 21 days after electroporation *in vivo* at P1 with *Gfp* or *Pou3f1*. Yellow arrows point to GFP+ cells that are DCX+/BRN3B-, whereas white arrows point to GFP+/DCX+/BRN3B+ cells. **(B-D)** Quantification of the proportion of GFP+/BRN3B+ (B), GFP+/DCX+ (C) or GFP+/DCX+/BRN3B+ (D) cells in *Gfp* or *Pou3f1* electroporated retinas (n = 7 animals per condition). **(F)** Representative images of retinal flat mounts from *Gfp*- or *Pou3f1*-electroporated retinas 60 days (D) after electroporation at P1. Images in the lower row are desaturated and inverted to better visualize axon fasciculation and projection to the optic nerve head (ONH) in the Pou3f1 condition. **(G)** Immunostaining for GFP and BRN3B on cross sections of Atoh7 knockout retinal explants 14 days after electroporation with *Gfp* or *Pou3f1* at P0. **(H)** Quantification of the percentage of GFP^+^/BRN3B+ cells in *Gfp* and *Pou3f1*-electroporated Atoh7 KO retinas.

We next asked whether POU3F1-induced RGC-L cells possess similar morphology and projection characteristics as endogenous RGCs. We thus dissected *Gfp* and *Pou3f1* electroporated retinas at 21 and 60 days and prepared retinal flatmounts to visualize axonal projections. While we did not observe projections emanating from the *Gfp* electroporated regions, as expected given that no RGCs are produced from P1 progenitors, we found several projections emerging from the *Pou3f1* electroporated regions at 21 days (Fig. S7), which was even more pronounced at 60 days (Fig. 7F). Remarkably, these *de novo* projections converged towards the optic nerve head (Fig. 7F), as do endogenous RGC axons. Therefore, although POU3F1-induced RGC-L cells do not express all RGC markers, they share many molecular and morphological features with *bona fide* RGCs.

The above data suggest that POU3F1 is sufficient for RGC-L cell production at a stage when ATOH7 is no longer expressed, and given our findings that POU3F1 represses *Atoh7*, we wondered whether POU3F1 might be part of the elusive ATOH7-independent RGC production pathway recently suggested (Brodie-Kommit *et al*., 2021). To test this hypothesis, we electroporated P1 *Atoh7*^lacZ/lacZ^ (KO) mice in vivo with *Pou3f1* or *Gfp* alone, collected the retinas 14 days later and stained for the RGC marker BRN3B. We found that expression of *Pou3f1* in *Atoh7* KO retinas is sufficient to induce production of BRN3B^+^ cells (Fig. 6F, G), which were never observed after expression of *Gfp* alone. These results show that POU3F1 is sufficient to promote the production of RGC-L cells in absence of ATOH7, suggesting that it is part of an ATOH7-independent RGC specification program.

## Discussion

Only a few factors involved in the development of the cRGC fate have been reported so far (Kuwajima *et al*., 2017; Pak *et al*., 2004), and how expression of these factors is regulated remains unclear. Our study now identifies POU3F1 as a master regulator of the cRGC transcriptional program (Fig. 8). We report that POU3F1 is expressed in developing cRGCs during mouse retinal development. Using loss- and gain-of-function approaches, we show that POU3F1 is required for cRGC development and promotes the cRGC fate choice when expressed in retinal progenitors. Moreover, RNA-seq and Cut&Run-Seq reveal that POU3F1 binds to and regulates the expression of genes involved in the iRGC/cRGC fate choice. A key finding is the identification of POU3F1 as a negative regulator of *Atoh7*, thereby leading to repression of *Zic2* and acquisition of the cRGC fate. Finally, we report that the sole expression of POU3F1 in postnatal progenitors is sufficient to produce cells sharing molecular and morphological features with RGCs, including axonal projection to the optic nerve head, and that POU3F1 expression can bypass the requirement for ATOH7 in RGC production, raising interesting possibilities for cell therapy approaches.

**Figure 8.**
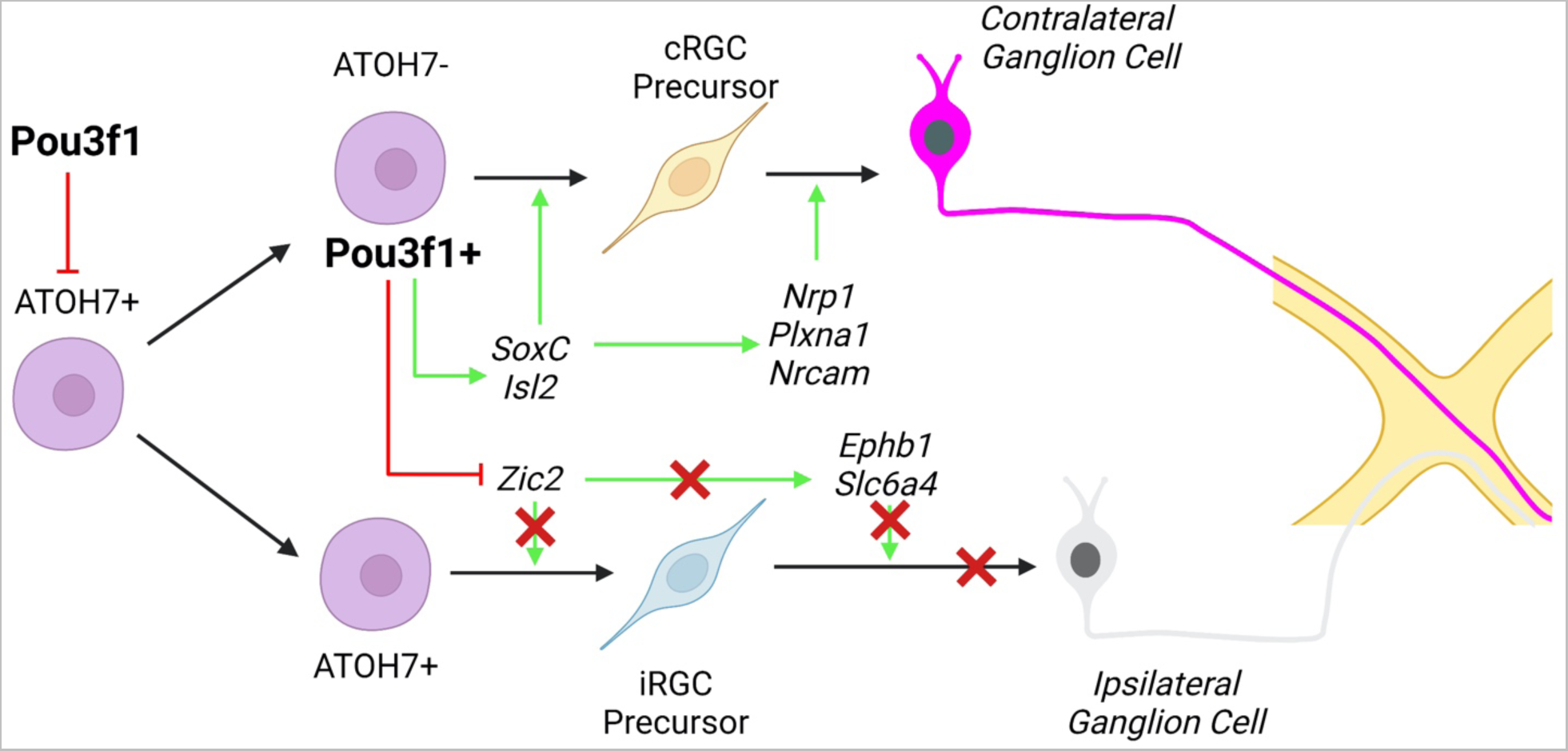
Proposed model of POU3F1 action in retinal development. POU3F1 represses *Atoh7* and activates *SoxC* and *Isl2*, promoting contralateral axonal determinants, while repressing *Zic2* and inhibiting ipsilateral determinants. This gene regulatory network leads to acquisition of the contralateral ganglion cell identity. Created with biorender.com.

### Competition between POU3F1 and ZIC2 contributes to establish the proper iRGC/cRGC ratio

While we did not find changes in *Zic2* expression levels by RNA-Seq, likely because E14.5 is too early and P1 + 48 hours is too late to pick up changes in *Zic2* transcript expression, we found that loss of POU3F1 leads to an expansion of the ZIC2^+^ cell population domain in both the VTA and the central retina at E17.5, and overexpression of *Pou3f1* in E14.5 explants reduces the number of ZIC2^+^ cells in the VTA. Additionally, our binding profile analysis of POU3F1 shows significant peaks on the *Zic2* promoter region. These results support the conclusion that POU3F1 represses ZIC2 in developing RGCs, thereby favoring cRGC fate acquisition. Interestingly, a recently published study showed that ZIC2 overexpression promotes the iRGC fate acquisition, which is accompanied by reduction of *Pou3f1* levels (Escalante *et al*., 2013). These observations suggest a model wherein ZIC2 and POU3F1 may compete to promote iRGCs and cRGCs, respectively. Of note, a similar rivalry has been observed in epiblast stem cells, where POU3F1 and ZIC2 promote the generation of the neural plate by repressing genes involved in mesoderm tissue maintenance (Iwafuchi-Doi *et al*., 2012; Zhu *et al*., 2014). Although POU3F1 and ZIC2 both promote the neural plate lineage, ChIP-Seq analysis revealed that their binding at target genes is mutually exclusive, in contrast with other neural plate inducing genes like *Otx2*, *Sox2* and *Pou5f1* (*Oct4*) (Matsuda *et al*., 2017). It remains unclear whether this kind of competition between POU3F1 and ZIC2 is a general logic operating in multiple systems, but some observations are consistent with this possibility. For example, misexpression of ZIC2 in the developing spinal cord is sufficient to induce ipsilateral identity of commissural neurons by downregulating genes that promote floor plate crossing, such as LHX2 and ROBO3 (Escalante *et al*., 2013). Intriguingly, POU3F1 expression has been reported at E11.5 in developing dI1-3, MNs, V2a and V3 interneurons and co-labels abundantly with LHX2 (Dasen et al., 2005; Delile et al., 2019; Francius et al., 2013), but its function in these cells remains unknown. Given that ZIC2 and POU3F1 are expressed in overlapping timepoints and areas in the developing spinal cord, it will be interesting to investigate their relationship on the establishment of axonal laterality. Our findings suggest that similar competition for binding at target genes may be involved.

### Selecting the contralateral identity among the pool of developing RGCs

We find that loss of POU3F1 does not affect the total number of RGCs produced but favors production of iRGC at the expense of cRGCs among the pool of differentiating RGCs. Using transcription profiling, we report that, in addition to repressing *Atoh7* and *Zic2*, POU3F1 generally promotes the expression of contralateral determinants. In *Pou3f1* KO retinas, we find that cRGC markers like *Igf1*, *Sema3e*, *Tfap2a*, and *Rgs4*, as well as the contralateral determinants *Sox4* (member of the *SoxC* family) and *Isl2* are downregulated (Kuwajima *et al*., 2017; Murcia-Belmonte and Erskine, 2019). Conversely, we find that misexpression of POU3F1 induces *Sox4/11* and the contralateral guidance receptors *Nrp1* and *PlexinA1.* In dissociated retinal cell cultures, SoxC triple KO (*Sox4/11/12*) have more than an 80% reduction in the number of cells labeled with contralateral RGC markers (Islet1/2), while in these same cultures, single inactivation of *Sox4* leads to approximately 40% decrease in iRGCs (Kuwajima *et al*., 2017), similar to what we find in the *Pou3f1* KO. SOXC was found to promote cRGCs by suppressing NOTCH signaling through binding at the *Hes5* promoter. Consistently, our data show that POU3F1 binds to a region upstream of *Sox4*, *Notch1* and *Hes5* promoters, and POU3F1 overexpression decreases both *Notch1* and *Hes5* transcript levels. These results suggest that POU3F1 contributes to suppressing NOTCH signaling to promote the cRGC fate, directly or indirectly via upregulation of *Sox4*.

ISL2 is another key determinant of the cRGC fate that is downregulated in the *Pou3f1* KO retina. *Isl2* null retinas display increased expression of the iRGC determinant *Zic2* and guidance receptor *EphB1* in the VTA, suggesting that ISL2 promotes the cRGC fate by repressing iRGC determinants (Pak *et al*., 2004). Similarly, we report here that loss of POU3F1 increases the number of ZIC2^+^ cells in the VTA and in the central retina. Moreover, overexpression of POU3F1 in the postnatal retina downregulates *Ephb1* levels in electroporated cells, suggesting a repressive role for POU3F1 on these major iRGC determinants. Consistently, the transcriptional changes observed in *Pou3f1* null retinas result in guidance abnormalities at the optic chiasm, with *Pou3f1* KO mice displaying increased number of RGC fibers on the ipsilateral side compared to their heterozygote littermates.

### Atoh7-independent generation of RGCs

Until recently, ATOH7 was believed to be the main driver of RGC development, and its downstream mechanisms have been extensively studied. Several studies showed that ATOH7 is necessary for RGC production during early retinogenesis, as *Atoh7*-deficient retinas lack 95% of RGCs (Brown *et al*., 2001). However, one recent study showed that co-expression of *Isl1* and *Brn3b* can rescue RGC production in *Atoh7* KO retinas (Wu et al., 2015). These results are consistent with previous findings demonstrating that both *Isl1* and *Brn3b* are direct downstream targets of ATOH7 (Gao *et al*., 2014; Mu et al., 2005). In general, the necessity of ATOH7 for RGC production has long been debated, as lineage studies revealed that only 50% of RGCs are part of the ATOH7 lineage (Brzezinski et al., 2012). A recent study provided clarifications on these questions and showed that ATOH7 is not necessary for RGC genesis per se, but rather essential for their survival (Brodie-Kommit *et al*., 2021). Indeed, preventing cell death in the *Atoh7* KO retina by inactivating BAX is sufficient to rescue RGC production to almost normal levels, although these RGCs failed to project to their appropriate targets (Brodie-Kommit *et al*., 2021). This study indicates that ATOH7-independent mechanisms exist to promote RGC genesis, but these remained unknown. Our data now show that POU3F1 is at least one of these factors controlling cRGC production independently of ATOH7. We observed that BRN3A/B are strongly induced by POU3F1 at embryonic stages and that BRN3B levels rise rapidly after ectopic POU3F1 expression in postnatal development when ATOH7 is no longer expressed. Moreover, we find that POU3F1 represses ATOH7, and yet is still able to induce RGC production. Finally, overexpression of POU3F1 in the postnatal ATOH7 KO retina is sufficient to promote RGC-L cell generation. It will be interesting in the future to determine whether POU3F1 expression in the embryonic *Atoh7* KO retina is able to rescue optic nerve development. While we report here that Pou3f1 expression in P1 retinas induce de novo production of RGC-L cells that project axons to the optic nerve head, they do not appear to enter the optic nerve. This may be because wildtype optic nerves are already filled with endogenous RGC axons, leaving no space for the POU3F1- induced RGC-L axons to grow. Expressing POU3F1 in embryonic *Atoh7* KO retinas, which have an optic tract but almost no axons, should help test this idea.

### Promoting axonogenesis with POU3F1: a candidate for regenerative therapies?

Initial studies of POU3F1 revealed its role in Schwann cell differentiation and the onset of myelination in the PNS (Blanchard et al., 1996; Jaegle *et al*., 2003). Recent work has demonstrated that it is also involved in promoting the neuronal fate in embryonic stem cells (ESCs) (Zhu *et al*., 2014). In this same study, and like the results presented here, GO analysis of *Pou3f1* overexpression in ESCs revealed that the most bound and expressed genes are associated with neuron development, neuron-projection morphogenesis and axonogenesis (Zhu *et al*., 2014). Consistently, rapid axon formation is robustly induced and consistently observed in our gain of function assays, hallmarked by the upregulation of several axon growth promoting (*Alcam, Slitrk1/2, Eomes and Dcx*) and guidance genes *(Dcc, Robo1/2, PlexinA1 and Nrp1*). Moreover, such potent *de novo* formation, self-organization and axonal projection of RGC-L cells from postnatal progenitors is, to our knowledge, unprecedented and suggest a possible role for POU3F1 in stimulating axonogenesis, which could be of interest for translational work aiming at regenerating the optic nerve. It remains unclear, however, whether the ectopically produced RGC-L cells are functional, as they fail to enter the optic nerve. As discussed above, this is possibly due to the optic nerve being saturated by endogenous RGC axons. In addition to exploring whether POU3F1 can induce axonal growth in the *Atoh7* KO optic nerves, it will be interesting to determine whether POU3F1 can stimulate axonal regeneration in the context of RGC degeneration. A recent study showed that expression of the OSK factors (*Oct4*, *Sox2* and *Klf4*) in axotomized RGCs can promote regeneration and restore visual functions (Lu et al., 2020). This regenerative phenotype required the DNA-demethylation activity of the Tet methylcytosine dioxygenase 2 (TET2). Interestingly, we report here that *Tet2* is upregulated by Pou3f1 (Fig. S5A, B), suggesting that POU3F1 may promote axonal growth in a TET2-dependent manner (Lu *et al*., 2020).

In conclusion, this work uncovers POU3F1 as a master regulator of cRGC production, thereby controlling the proper iRGC/cRGC ratio. These findings have implications for our understanding of the molecular mechanisms underlying the emergence of binocular vision in mammals and open new possibilities for the development of regenerative therapies.

## Material and Methods

### Animals

All listed experiments were performed in accordance with the Canadian Council on Animal guidelines. Pou3f1-ßGeo (C57/Bl6 x Sv129) was previously generated, characterized and kindly obtained from Dies Meijer (University of Edinburgh, Edinburgh, UK) (Jaegle *et al*., 2003; Jaegle et al., 1996). Pou3f1-ßGeo pregnant females were sacrificed at different embryonic timepoints depending on the assay followed by immediate embryo extraction. Given that a high percentage of Pou3f1-ßGeo KO die perinatally, all crosses were performed using Pou3f1-ßGeo HET. The Pou3f1-ßGeo allele was genotyped by PCR following extraction. Sert-Cre (5’HTT-Cre, ET33) mice were obtained from the Jackson Laboratory and crossed with a R26-tdtomato reporter line(Gong et al., 2007). Time-course, gain-of-function RNA-seq and Cut&Run experiments were all carried out in a wildtype (WT) C57/Bl6 background. *Ex vivo* and *In vivo* gain-of-function assays were performed in a CD1 background (*Mus Musculus*, Charles River). *Atoh7* KO mice (*Atoh7^tm1Gla^*) have been previously characterized and were obtained from the MMRRC at UC Davis, California, USA (Brown *et al*., 2001; Miesfeld *et al*., 2018).

### Tissue collection and Immunofluorescence

All female pregnancies were timed, and the day of fertilization was set at day 0.5 (E0.5). Pregnant females were sacrificed at E13.5, E14.5, E15.5, E16.5 or E17.5 by cervical dislocation and embryos were extracted afterwards. Embryos were decapitated using dissection scissors and heads were washed once in sterile PBS (Gibco^TM^; Cat-Nr.: 14190144). Heads were fixed in freshly prepared 4% PFA/PBS for 15 minutes to 1 hour at room temperature (RT) followed by another 5-minute PBS wash. Finally, heads were immersed for 2 hours to overnight (O/N) in a 20% Sucrose/PBS solution at 4°C. Postnatal day (P) mice were sacrificed either by decapitation (P1-14) or CO2 asphyxiation followed by cervical dislocation (>P14). Eyes were extracted and the cornea was punctured before immersion in 4% PFA O/N at 4°C, followed by cryoprotection with 20% Sucrose. After sucrose cryoprotection, heads or eyes were placed in a large cryomold and embedded in 20% Sucrose:OCT (1:1 ratio) solution (OCT, Sakura, 4583) before freezing using liquid nitrogen. Head tissue was sectioned at 25 µm and eyes at 18 µm using a cryostat (Leica CM3050S) with an air temperature setting of -20°C onto microscope glass slides (Fisher Scientific).

For immunofluorescence, slides were dipped briefly in Phosphate-buffered saline (PBS) to remove excess OCT and left to dry for 15 minutes before mounting them on Sequenza racks. Sections were blocked with PBS containing 3% Bovine Serum Albumin (BSA) (Thermo Fischer) and 0.5% Triton-X for 45 minutes before adding primary antibodies. The following antibodies were used for this study; anti-POU3F1 rabbit (1:100, Genetex, GTX134063), anti-POU3F1 mouse (1:100, EMD Millipore Sigma, MABN738), anti-POU3F1 rabbit #1909 and #1894 (both 1:100, gift from Dr. Meijer Dies, University of Edinburgh, Edinburgh, UK), anti-ZIC2 rabbit (1:10000, gift from Dr. Carol Mason, Columbia University), anti-ATOH7 rabbit (1:100, Novus Biologicals, NBP1-88639), anti- BRN3A guinea-pig (1:10000, Synaptic Systems, 411004), anti-BRN3B goat (1:500, Santa Cruz, discontinued), anti-BRN3B goat (1:200, Rockland Scientific, 600-101-MJ0), anti-ISL1 mouse (1:50, DHSB, 40.2D6), anti-PAX6 rabbit (1:200, EMD Millipore Sigma, AB22337), anti-LHX2 rabbit (1:250, Thermo Fisher Scientific, PA5-78287), anti-CHX10 sheep (1:500, Exalpha Biologicals, X1180P), anti-BIII-Tubulin mouse (1:1000, Covance, MMS-435P), anti-DCX guinea-pig (1:400, EMD Millipore Sigma, AB2253) and Click-iT EdU reaction kit, AlexaFluor-647 (Thermo Fisher Scientific, C10340). Sections were incubated with primary antibodies overnight at RT and washed three times with PBS (5 minutes per wash) before adding the secondary antibodies for 1 hour at RT. At the end of immunostaining, sections were washed again three times with the third wash containing Hoechst (1:10000, Invitrogen, 33342). Slides were cover-slipped with Mowiol (Calbiochem.) and left to dry in darkness overnight before imaging.

### Electroporations, retinal explant culture and EdU birthdating

*Ex vivo, in vivo* electroporations and retinal explant cultures were performed as previously described(Javed et al., 2020). For *ex vivo* electroporations, P1 eyes were injected sub- retinally with 2ug of DNA resuspended in 1ul of water containing 0.5% of fast-green and underwent 5 pulses at 50V, 50ms. Explants were fixed at different timepoints depending on the assay. For *ex vivo* birth dating experiments, culture medium containing 5µM EdU was added to the well for 2 hours before fixation or for 1 hour after electroporation. For *in vivo* electroporations, whole heads received 5 pulses at 80V, 50ms each. Mice were sacrificed either at 2, 14, 21 days or 6 weeks after electroporation. Tissues were extracted and processed as stated above. For *in vivo* EdU birth dating, we injected pregnant females intraperitoneally at stage E17.5 with 50μg of EdU per gram of body weight followed by a 2-hour pulse before sacrifice. We extracted the embryos, collected the eyes and fixed for 10 minutes at RT in 4% PFA. Eyes were cryoprotected, as previously described and embedded in OCT prior to snap-freezing. Cryosections were stained for Atoh7, Zic2 and with a click-it EdU cell proliferation kit (Thermo Fisher, C10638) was used after secondary antibody incubation by following manufacturer’s recommendations.

### Plasmids

*Pou3f1* and *Atoh7* cDNA were obtained as PCR products using the following primers: For pCIG-Pou3f1: 5’- GGGGACAAGTTTGTACAAAAAAGCAGGCTTAATGGCCACCACCGCGCAGTATCTGC-3’ and 5’- GGGGACCACTTTGTACAAGAAAGCTGGGTTTCACTGCACAGAGCCGGGCAGTGTGT-3’; for pCig-Atoh7: 5’-CTCAAGCTTCGAATTATGAAGTCGGCCTGCAAACC-3’, and 5’-GTCGACTGCAGAATTTTAGCTGGCCATGGGGAAGG-3’. All cDNA fragments were cloned into a pCIG2 (pCAG-IRES-GFP) backbone using the InFusion® HD Cloning Kit (TakaraBio, 639650). The *Zic2* plasmid (pCAG-*Zic2*) was a gift from Dr. Eloisa Herrera (Instituto de Neurosciencias, Alicante, Spain).

### ß-Gal staining and cell sorting

Retinas were extracted, rapidly isolated, and placed into individual 1.5ml DNA LoBind® tubes (Eppendorf, 0030122348) containing 50µl PBS and kept on ice until dissociation. Retinas were dissociated with 40 units of Papain (Worthington, LS003126) and centrifuged at 300g for 5 minutes. For sorting of ßGal+ cells from Atoh7^lacZ/lacZ^ or Atoh7^lacZ/+^, the pellet was resuspended in staining media and ßGal activity was detected using FluoReporter^TM^ *lacZ* Flow Cytometry Kit (Thermo Fisher Scientific, F1930) according to manufacturer’s guidelines. For sorting of GFP expressing cells, no staining was performed. The resulting pellet was resuspended in 80µl of ice-cold PBS and cells isolated with a BD FACSAria^TM^ III Cell Sorter using the 488-laser as a gating channel. Sorted ßGal+ or GFP^+^ cells were collected in 1.5ml DNA LoBind® tubes placed at 4°C containing 400µl RLT buffer (Qiagen, 79216) with 1% β-Mercaptoethanol (Millipore Sigma, M6250) for RNA-extraction.

### RNA extraction

RLT buffer containing either sorted cells or dissociated retinas were vortexed with maximum strength for 3min to induce proper cell lysis and maximal RNA release. RNA was extracted using the RNA micro kit (Qiagen, 74004) according to manufacturer’s protocol if the number of FACS collected cells was inferior to a total of 500 000 cells per condition. Total RNA was eluted in 14µl RNAse free water and stored at -80°C until further use.

### RNASeq and analysis

Directional mRNA libraries were obtained using the Illumina TruSeq mRNA kit. RNA samples were then converted to cDNA libraries by reverse transcription, followed by a PCR amplification and subsequent sequencing on Illumina HiSeq2500. Paired sequencing results were analyzed on the open access platform Galaxy (usegalaxy.org) using the functions *Salmonquant* followed by *DESeq2* to monitor differential gene expression and were aligned on the mm10 (*Mus musculus*) genome. *GOrilla (Gene Ontology enrichment analysis and visualization tool)* was used to perform the GO analysis of both significantly up- and down-regulated gene fractions(Eden et al., 2007; Eden et al., 2009)

### Cut&Run Seq

For each condition 2ug of rabbit anti-POU3F1 #1909, 2ug of rabbit anti-POU3F1 #1894 and 2ug of rabbit isotype control IgG (Thermofisher Scientific, 02-6102) antibodies were added to the samples and incubated at 4°C overnight. Sequencing results were analyzed with the open access platform Galaxy using the following commands *Trimmomatic, Bowtie2, MACS2 callpeak (FDR = 0.05*) followed by *bamCoverage*. The BED file from the *MACS2 callpeak* analysis was used to perform the GO analysis using GREAT(McLean *et al*., 2010).

### DiI tracing

Embryos were extracted at E17.5 and heads were washed three times in PBS before transferring them in individual wells of a 24-well plate. 2µl of DiI dissolved in DMSO (2.5μg/μl) was injected under the microscope into the right eye of each head followed by an overnight incubation in 4% PFA at 37°C, heads were then tightly sealed with paraffin for a 10-day incubation on a light shaker at 37 °C. After incubation, heads were washed 3x in PBS and transferred to 20% sucrose and overnight at 4 °C before being embedded in OCT and sectioned at 30µm in coronal orientation.

### Imaging and quantification

Immunofluorescence and DiI tracing were imaged using a Leica SP8 Confocal Microscope. For DiI tracing at the optic nerve and caudal chiasm, images were acquired with a 5X lens and 16-bit resolution. Laser intensity was kept consistent for all sections.

Average intensity quantifications of contra- and ipsilateral tracts were selected and measured on ImageJ using “mean gray value”. The average intensity of the selected area was background corrected by subtracting the background average intensity by moving the selection to an area adjacent to the optic tract. I-ratio was determined with the following calculation:

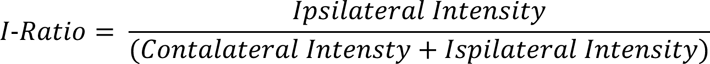

For image analysis and cell quantifications, a FFT Bandpass filter was applied to limit background selections on the SP8-acquired images. Positive cells were selected by a manually defined fluorescence intensity threshold whose settings was kept the same for all samples and condition. Cells were then automatically counted using the “Analyze Particle” plugin in ImageJ. For electroporation, ATOH7 and ZIC2 staining, GFP and all cell markers were counted manually except for DAPI. Population percentages were calculated by dividing the total cell number of each cellular marker by DAPI.

### Statistical and Quantitative analysis

All figures were created using the software GraphPad Prism ver.9. For comparisons between two conditions, unpaired, t-tests with Mann-Whitney post tests were performed with a significant cut-off at p<0.05 unless indicated otherwise. One-way ANOVA and Tukey’s post-hoc test was performed for multiple comparisons. All error bars represent SEM and asterisk are indicated as the following: p* < 0.05, p** < 0.01, p*** < 0.001, p****< 0.0001. No asterisks between conditions meant p > 0.05. For the RNA-Seq, all values under p < 0.05 were considered as significant hits.

## Acknowledgements

We thank Seth Blackshaw for comments on the manuscript and the IRCM core facilities for technical assistance, in particular: Jessica Barthe for mouse colony management and Dominic Filion for help with microscopy and image analysis. We are also grateful to all members of the Cayouette lab, past and present, for ongoing suggestions on this work. This study was supported by grants from the Canadian Institutes of Health Research (FDN-159936) to M.C. A.J. and C.B-P. received studentships from Fonds de recherche du Québec Santé (FRQS). M.C. is an Emeritus Research Scholar from the Fonds de recherche du Québec Santé (FRQS) and holds the Gaëtane and Roland Pillenière Chair in Retina Biology from the IRCM Foundation.

## References

1. Andermatt, I., Wilson, N.H., Bergmann, T., Mauti, O., Gesemann, M., Sockanathan, S., and Stoeckli, E.T. (2014). Semaphorin 6B acts as a receptor in post-crossing commissural axon guidance. Development 141, 3709–3720. 10.1242/dev.112185.

2. Bermingham, J.R., Jr., Scherer, S.S., O’Connell, S., Arroyo, E., Kalla, K.A., Powell, F.L., and Rosenfeld, M.G. (1996). Tst-1/Oct-6/SCIP regulates a unique step in peripheral myelination and is required for normal respiration. Genes Dev 10, 1751–1762. 10.1101/gad.10.14.1751.

3. Blanchard, A.D., Sinanan, A., Parmantier, E., Zwart, R., Broos, L., Meijer, D., Meier, C., Jessen, K.R., and Mirsky, R. (1996). Oct-6 (SCIP/Tst-1) is expressed in Schwann cell precursors, embryonic Schwann cells, and postnatal myelinating Schwann cells: comparison with Oct-1, Krox-20, and Pax-3. J Neurosci Res 46, 630-640. 10.1002/(SICI)1097-4547(19961201)46:5<630::AID-JNR11>3.0.CO;2-0.

4. Brodie-Kommit, J., Clark, B.S., Shi, Q., Shiau, F., Kim, D.W., Langel, J., Sheely, C., Ruzycki, P.A., Fries, M., Javed, A., et al. (2021). Atoh7-independent specification of retinal ganglion cell identity. Sci Adv 7. 10.1126/sciadv.abe4983.

5. Brown, N.L., Patel, S., Brzezinski, J., and Glaser, T. (2001). Math5 is required for retinal ganglion cell and optic nerve formation. Development 128, 2497–2508.

6. Brzezinski, J.A.t., Prasov, L., and Glaser, T. (2012). Math5 defines the ganglion cell competence state in a subpopulation of retinal progenitor cells exiting the cell cycle. Dev Biol 365, 395–413. 10.1016/j.ydbio.2012.03.006.

7. Clark, B.S., Stein-O’Brien, G.L., Shiau, F., Cannon, G.H., Davis-Marcisak, E., Sherman, T., Santiago, C.P., Hoang, T.V., Rajaii, F., James-Esposito, R.E., et al. (2019). Single-Cell RNA-Seq Analysis of Retinal Development Identifies NFI Factors as Regulating Mitotic Exit and Late-Born Cell Specification. Neuron 102, 1111–1126 e1115. 10.1016/j.neuron.2019.04.010.

8. Dasen, J.S., Tice, B.C., Brenner-Morton, S., and Jessell, T.M. (2005). A Hox regulatory network establishes motor neuron pool identity and target-muscle connectivity. Cell 123, 477–491. 10.1016/j.cell.2005.09.009.

9. Deiner, M.S., Kennedy, T.E., Fazeli, A., Serafini, T., Tessier-Lavigne, M., and Sretavan, D.W. (1997). Netrin-1 and DCC mediate axon guidance locally at the optic disc: loss of function leads to optic nerve hypoplasia. Neuron 19, 575–589. 10.1016/s0896-6273(00)80373-6.

10. Delile, J., Rayon, T., Melchionda, M., Edwards, A., Briscoe, J., and Sagner, A. (2019). Single cell transcriptomics reveals spatial and temporal dynamics of gene expression in the developing mouse spinal cord. Development 146. 10.1242/dev.173807.

11. Ding, Q., Joshi, P.S., Xie, Z.H., Xiang, M., and Gan, L. (2012). BARHL2 transcription factor regulates the ipsilateral/contralateral subtype divergence in postmitotic dI1 neurons of the developing spinal cord. Proc Natl Acad Sci U S A 109, 1566–1571. 10.1073/pnas.1112392109.

12. Drayson, L.E., and Triplett, J.W. (2019). A Chrnb3-Cre BAC transgenic mouse line for manipulation of gene expression in retinal ganglion cells. Genesis 57, e23305. 10.1002/dvg.23305.

13. Duan, X., Krishnaswamy, A., Laboulaye, M.A., Liu, J., Peng, Y.R., Yamagata, M., Toma, K., and Sanes, J.R. (2018). Cadherin Combinations Recruit Dendrites of Distinct Retinal Neurons to a Shared Interneuronal Scaffold. Neuron 99, 1145–1154 e1146. 10.1016/j.neuron.2018.08.019.

14. Eden, E., Lipson, D., Yogev, S., and Yakhini, Z. (2007). Discovering motifs in ranked lists of DNA sequences. PLoS Comput Biol 3, e39. 10.1371/journal.pcbi.0030039.

15. Eden, E., Navon, R., Steinfeld, I., Lipson, D., and Yakhini, Z. (2009). GOrilla: a tool for discovery and visualization of enriched GO terms in ranked gene lists. BMC Bioinformatics 10, 48. 10.1186/1471-2105-10-48.

16. Erskine, L., Reijntjes, S., Pratt, T., Denti, L., Schwarz, Q., Vieira, J.M., Alakakone, B., Shewan, D., and Ruhrberg, C. (2011). VEGF signaling through neuropilin 1 guides commissural axon crossing at the optic chiasm. Neuron 70, 951–965. 10.1016/j.neuron.2011.02.052.

17. Erskine, L., Williams, S.E., Brose, K., Kidd, T., Rachel, R.A., Goodman, C.S., Tessier-Lavigne, M., and Mason, C.A. (2000). Retinal ganglion cell axon guidance in the mouse optic chiasm: expression and function of robos and slits. J Neurosci 20, 4975–4982.

18. Escalante, A., Murillo, B., Morenilla-Palao, C., Klar, A., and Herrera, E. (2013). Zic2- dependent axon midline avoidance controls the formation of major ipsilateral tracts in the CNS. Neuron 80, 1392–1406. 10.1016/j.neuron.2013.10.007.

19. Francius, C., Harris, A., Rucchin, V., Hendricks, T.J., Stam, F.J., Barber, M., Kurek, D., Grosveld, F.G., Pierani, A., Goulding, M., and Clotman, F. (2013). Identification of multiple subsets of ventral interneurons and differential distribution along the rostrocaudal axis of the developing spinal cord. PLoS One 8, e70325 10.1371/journal.pone.0070325.

20. Fruttiger, M., Calver, A.R., Kruger, W.H., Mudhar, H.S., Michalovich, D., Takakura, N., Nishikawa, S., and Richardson, W.D. (1996). PDGF mediates a neuron-astrocyte interaction in the developing retina. Neuron 17, 1117–1131. 10.1016/s0896-6273(00)80244-5.

21. Gao, Z., Mao, C.A., Pan, P., Mu, X., and Klein, W.H. (2014). Transcriptome of Atoh7 retinal progenitor cells identifies new Atoh7-dependent regulatory genes for retinal ganglion cell formation. Dev Neurobiol 74, 1123–1140. 10.1002/dneu.22188.

22. Garcia-Frigola, C., Carreres, M.I., Vegar, C., Mason, C., and Herrera, E. (2008). Zic2 promotes axonal divergence at the optic chiasm midline by EphB1-dependent and - independent mechanisms. Development 135, 1833–1841. 10.1242/dev.020693.

23. Garcia-Frigola, C., and Herrera, E. (2010). Zic2 regulates the expression of Sert to modulate eye-specific refinement at the visual targets. EMBO J 29, 3170–3183. 10.1038/emboj.2010.172.

24. Gong, S., Doughty, M., Harbaugh, C.R., Cummins, A., Hatten, M.E., Heintz, N., and Gerfen, C.R. (2007). Targeting Cre recombinase to specific neuron populations with bacterial artificial chromosome constructs. J Neurosci 27, 9817–9823. 10.1523/JNEUROSCI.2707-07.2007.

25. Herrera, E., Brown, L., Aruga, J., Rachel, R.A., Dolen, G., Mikoshiba, K., Brown, S., and Mason, C.A. (2003). Zic2 patterns binocular vision by specifying the uncrossed retinal projection. Cell 114, 545–557. 10.1016/s0092-8674(03)00684-6.

26. Iwafuchi-Doi, M., Matsuda, K., Murakami, K., Niwa, H., Tesar, P.J., Aruga, J., Matsuo, I., and Kondoh, H. (2012). Transcriptional regulatory networks in epiblast cells and during anterior neural plate development as modeled in epiblast stem cells. Development 139, 3926–3937. 10.1242/dev.085936.

27. Jaegle, M., Ghazvini, M., Mandemakers, W., Piirsoo, M., Driegen, S., Levavasseur, F., Raghoenath, S., Grosveld, F., and Meijer, D. (2003). The POU proteins Brn-2 and Oct-6 share important functions in Schwann cell development. Genes Dev 17, 1380–1391. 10.1101/gad.258203.

28. Jaegle, M., Mandemakers, W., Broos, L., Zwart, R., Karis, A., Visser, P., Grosveld, F., and Meijer, D. (1996). The POU factor Oct-6 and Schwann cell differentiation. Science 273, 507–510. 10.1126/science.273.5274.507.

29. Javed, A., Mattar, P., Lu, S., Kruczek, K., Kloc, M., Gonzalez-Cordero, A., Bremner, R., Ali, R.R., and Cayouette, M. (2020). Pou2f1 and Pou2f2 cooperate to control the timing of cone photoreceptor production in the developing mouse retina. Development 147. 10.1242/dev.188730.

30. Kolodkin, A.L., and Tessier-Lavigne, M. (2011). Mechanisms and molecules of neuronal wiring: a primer. Cold Spring Harb Perspect Biol 3. 10.1101/cshperspect.a001727.

31. Kuwajima, T., Soares, C.A., Sitko, A.A., Lefebvre, V., and Mason, C. (2017). SoxC Transcription Factors Promote Contralateral Retinal Ganglion Cell Differentiation and Axon Guidance in the Mouse Visual System. Neuron 93, 1110–1125 e1115. 10.1016/j.neuron.2017.01.029.

32. Kuwajima, T., Yoshida, Y., Takegahara, N., Petros, T.J., Kumanogoh, A., Jessell, T.M., Sakurai, T., and Mason, C. (2012). Optic chiasm presentation of Semaphorin6D in the context of Plexin-A1 and Nr-CAM promotes retinal axon midline crossing. Neuron 74, 676–690. 10.1016/j.neuron.2012.03.025.

33. Lee, R., Petros, T.J., and Mason, C.A. (2008). Zic2 regulates retinal ganglion cell axon avoidance of ephrinB2 through inducing expression of the guidance receptor EphB1. J Neurosci 28, 5910–5919. 10.1523/JNEUROSCI.0632-08.2008.

34. Lu, Y., Brommer, B., Tian, X., Krishnan, A., Meer, M., Wang, C., Vera, D.L., Zeng, Q., Yu, D., Bonkowski, M.S., et al. (2020). Reprogramming to recover youthful epigenetic information and restore vision. Nature 588, 124–129. 10.1038/s41586-020-2975-4.

35. Manabe, S., Kashii, S., Honda, Y., Yamamoto, R., Katsuki, H., and Akaike, A. (2002). Quantification of axotomized ganglion cell death by explant culture of the rat retina. Neurosci Lett 334, 33–36. 10.1016/s0304-3940(02)01047-9.

36. Mao, C.A., Kiyama, T., Pan, P., Furuta, Y., Hadjantonakis, A.K., and Klein, W.H. (2008). Eomesodermin, a target gene of Pou4f2, is required for retinal ganglion cell and optic nerve development in the mouse. Development 135, 271–280. 10.1242/dev.009688.

37. Matsuda, K., Mikami, T., Oki, S., Iida, H., Andrabi, M., Boss, J.M., Yamaguchi, K., Shigenobu, S., and Kondoh, H. (2017). ChIP-seq analysis of genomic binding regions of five major transcription factors highlights a central role for ZIC2 in the mouse epiblast stem cell gene regulatory network. Development 144, 1948–1958. 10.1242/dev.143479.

38. McLean, C.Y., Bristor, D., Hiller, M., Clarke, S.L., Schaar, B.T., Lowe, C.B., Wenger, A.M., and Bejerano, G. (2010). GREAT improves functional interpretation of cis-regulatory regions. Nat Biotechnol 28, 495-501. 10.1038/nbt.1630.

39. McNeill, E.M., Roos, K.P., Moechars, D., and Clagett-Dame, M. (2010). Nav2 is necessary for cranial nerve development and blood pressure regulation. Neural Dev 5, 6. 10.1186/1749-8104-5-6.

40. Miesfeld, J.B., Glaser, T., and Brown, N.L. (2018). The dynamics of native Atoh7 protein expression during mouse retinal histogenesis, revealed with a new antibody. Gene Expr Patterns 27, 114–121. 10.1016/j.gep.2017.11.006.

41. Mu, X., Fu, X., Sun, H., Beremand, P.D., Thomas, T.L., and Klein, W.H. (2005). A gene network downstream of transcription factor Math5 regulates retinal progenitor cell competence and ganglion cell fate. Dev Biol 280, 467–481. 10.1016/j.ydbio.2005.01.028.

42. Murcia-Belmonte, V., and Erskine, L. (2019). Wiring the Binocular Visual Pathways. Int J Mol Sci 20. 10.3390/ijms20133282.

43. Nityananda, V., and Read, J.C.A. (2017). Stereopsis in animals: evolution, function and mechanisms. J Exp Biol 220, 2502–2512. 10.1242/jeb.143883.

44. Pak, W., Hindges, R., Lim, Y.S., Pfaff, S.L., and O’Leary, D.D. (2004). Magnitude of binocular vision controlled by islet-2 repression of a genetic program that specifies laterality of retinal axon pathfinding. Cell 119, 567–578. 10.1016/j.cell.2004.10.026.

45. Panza, P., Sitko, A.A., Maischein, H.M., Koch, I., Flotenmeyer, M., Wright, G.J., Mandai, K., Mason, C.A., and Sollner, C. (2015). The LRR receptor Islr2 is required for retinal axon routing at the vertebrate optic chiasm. Neural Dev 10, 23. 10.1186/s13064-015-0050-x.

46. Patterson, R., and Martin, W.L. (1992). Human stereopsis. Hum Factors 34, 669–692. 10.1177/001872089203400603.

47. Peng, J., Fabre, P.J., Dolique, T., Swikert, S.M., Kermasson, L., Shimogori, T., and Charron, F. (2018). Sonic Hedgehog Is a Remotely Produced Cue that Controls Axon Guidance Trans-axonally at a Midline Choice Point. Neuron 97, 326–340 e324. 10.1016/j.neuron.2017.12.028.

48. Quina, L.A., Pak, W., Lanier, J., Banwait, P., Gratwick, K., Liu, Y., Velasquez, T., O’Leary, D.D., Goulding, M., and Turner, E.E. (2005). Brn3a-expressing retinal ganglion cells project specifically to thalamocortical and collicular visual pathways. J Neurosci 25, 11595–11604. 10.1523/JNEUROSCI.2837-05.2005.

49. Ringstedt, T., Braisted, J.E., Brose, K., Kidd, T., Goodman, C., Tessier-Lavigne, M., and O’Leary, D.D. (2000). Slit inhibition of retinal axon growth and its role in retinal axon pathfinding and innervation patterns in the diencephalon. J Neurosci 20, 4983–4991.

50. Thelen, K., Maier, B., Faber, M., Albrecht, C., Fischer, P., and Pollerberg, G.E. (2012). Translation of the cell adhesion molecule ALCAM in axonal growth cones - regulation and functional importance. J Cell Sci 125, 1003–1014. 10.1242/jcs.096149.

51. Wakabayashi, T., Kosaka, J., Mori, T., Takamori, Y., and Yamada, H. (2008). Doublecortin expression continues into adulthood in horizontal cells in the rat retina. Neurosci Lett 442, 249–252. 10.1016/j.neulet.2008.07.030.

52. Wang, Q., Marcucci, F., Cerullo, I., and Mason, C. (2016). Ipsilateral and Contralateral Retinal Ganglion Cells Express Distinct Genes during Decussation at the Optic Chiasm. eNeuro 3. 10.1523/ENEURO.0169-16.2016.

53. Wu, F., Kaczynski, T.J., Sethuramanujam, S., Li, R., Jain, V., Slaughter, M., and Mu, X. (2015). Two transcription factors, Pou4f2 and Isl1, are sufficient to specify the retinal ganglion cell fate. Proc Natl Acad Sci U S A 112, E1559-1568. 10.1073/pnas.1421535112.

54. Zhu, Q., Song, L., Peng, G., Sun, N., Chen, J., Zhang, T., Sheng, N., Tang, W., Qian, C., Qiao, Y., et al. (2014). The transcription factor Pou3f1 promotes neural fate commitment via activation of neural lineage genes and inhibition of external signaling pathways. Elife 3. 10.7554/eLife.02224.

