## Supplemental Figures for "Pou3f1 orchestrates a gene regulatory network controlling contralateral retinogeniculate projections"

### SUPPLEMENTARY FIGURES

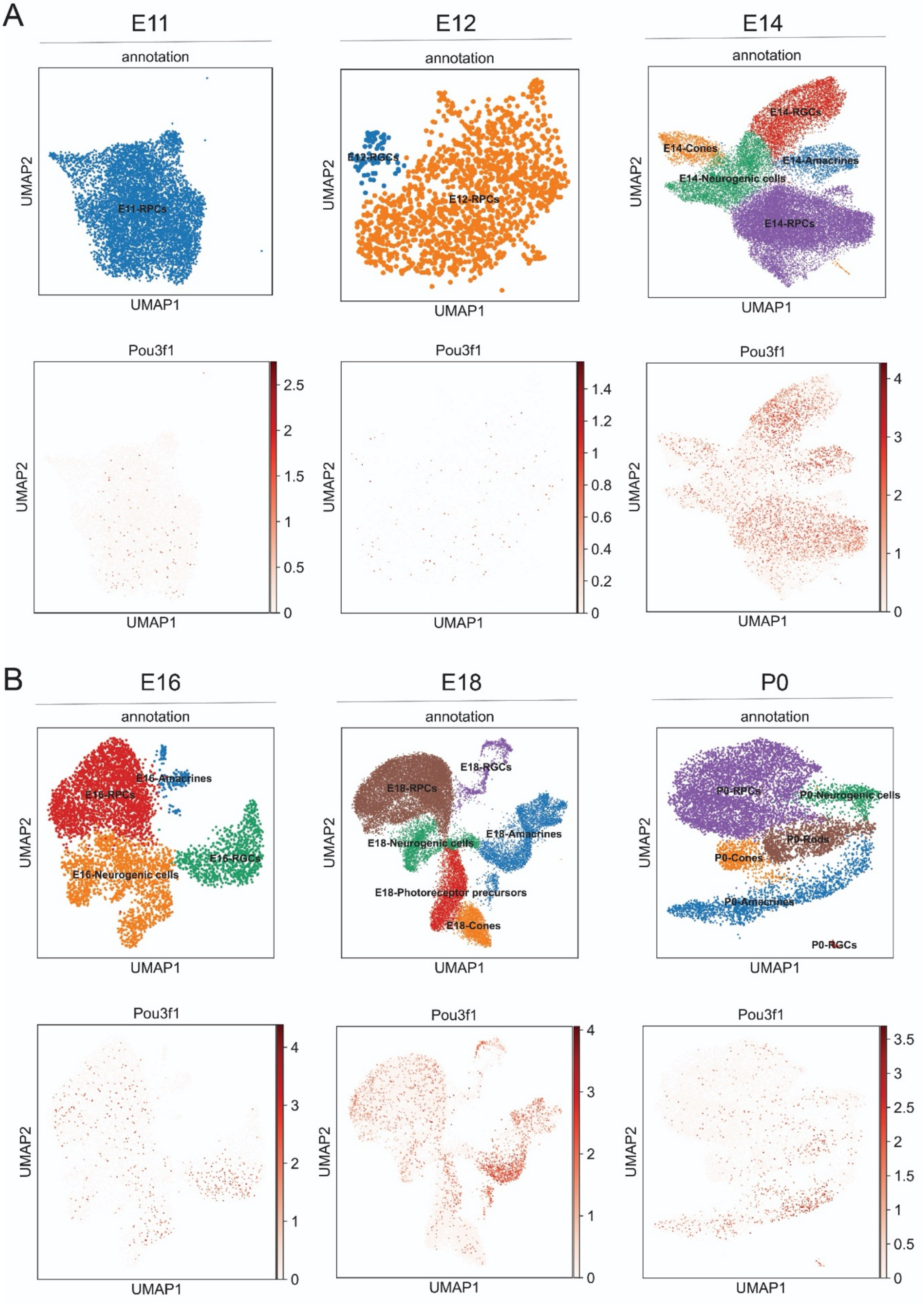

**Figure S1.** ***Pou3f1*** **expression in published scRNAseq datasets**

UMAP plots of annotated populations (top panels) and Pou3f1 expression (bottom panels) at different developmental stages. Data obtained from Clark et al., 2019 [1].

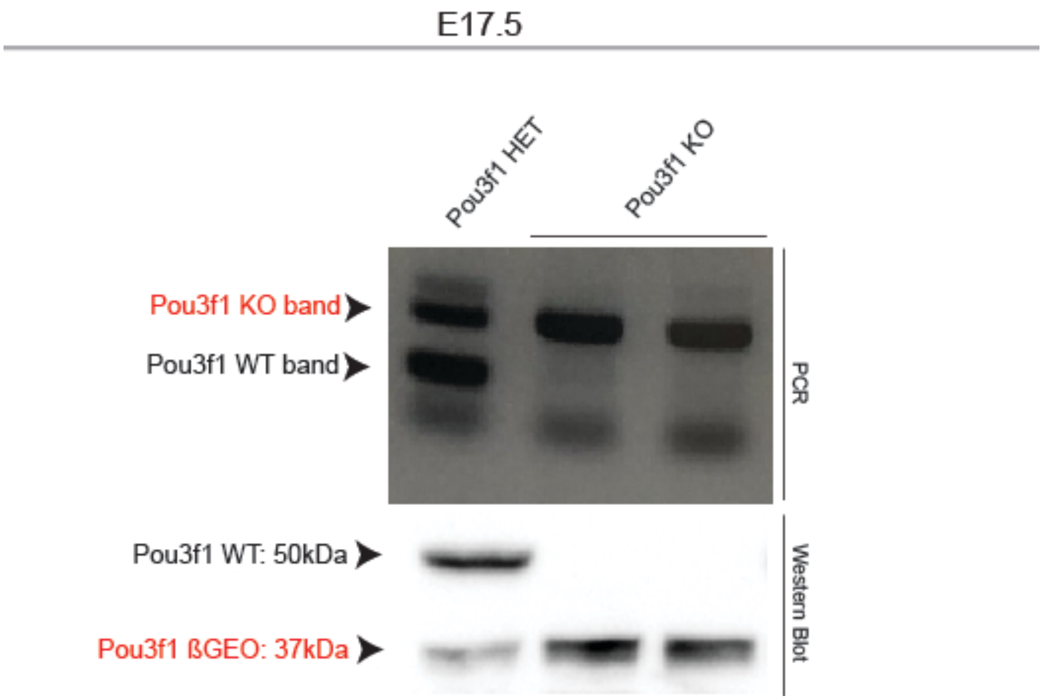


**Figure S2. Presence of a truncated POU3F1 protein in retinal extracts from *Pou3f1* KO mice.** PCR (Top) and Western Blot (Bottom) carried out on retinal extracts prepared from *Pou3f1* HET and *Pou3f1* KO mice. The truncated protein is observed in *Pou3f1* KO at 37kDa by Western blot.

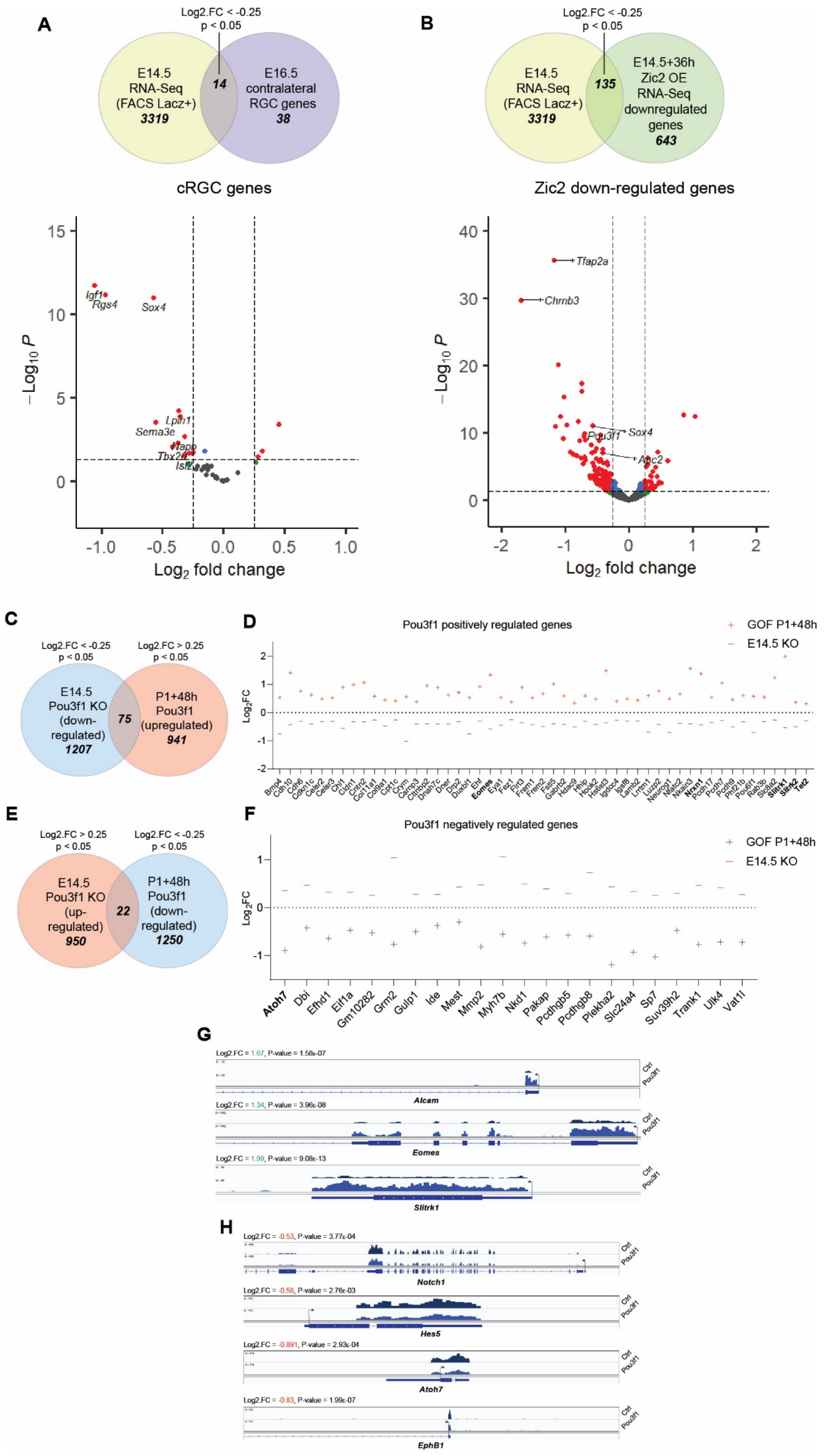

A
Log2.FC < -0.25
p < 0.05
E14.5
RNA-Seq
(FACS Lacz+)
3319
14
E16.5
contralateral
RGC genes
38
cRGC genes
B
Log2.FC < -0.25
p < 0.05
E14.5
RNA-Seq
(FACS Lacz+)
3319
135
E14.5+36h
Zic2 OE
RNA-Seq
downregulated
genes
643
Zic2 down-regulated genes
Log2 fold change
−Log10 P
Igf1
Rgs4
Sox4
Lpin1
Sema3e
Napp
Tbx20
Isl2
Tfap2a
Chrnb3
Pou3f1
Sox4
Apc2
C
Log2.FC < -0.25
p < 0.05
Log2.FC > 0.25
p < 0.05
E14.5
Pou3f1 KO
(down-
regulated)
1207
75
P1+48h
Pou3f1
(upregulated)
941
D
Pou3f1 positively regulated genes
GOF P1+48h
E14.5 KO
Log2FC
E
Log2.FC > 0.25
p < 0.05
Log2.FC < -0.25
p < 0.05
E14.5
Pou3f1 KO
(up-
regulated)
950
22
P1+48h
Pou3f1
(down-
regulated)
1250
F
Pou3f1 negatively regulated genes
GOF P1+48h
E14.5 KO
Log2FC
Atoh7
Dbi
Efhd1
Eif1a
Gm10282
Grm2
Gulp1
Ide
Mest
Mmp2
Myh7b
Nkd1
Pakap
Pcdhgb5
Pcdhgb8
Plekha2
Slc24a4
Sp7
Suv39h2
Trank1
Ulk4
Vat1l
G
Log2.FC = 1.67, P-value = 1.58e-07
Alcam
Log2.FC = 1.34, P-value = 3.96e-08
Eomes
Log2.FC = 1.99, P-value = 9.08e-13
Slitrk1
Ctrl
Pou3f1
H
Log2.FC = -0.53, P-value = 3.77e-04
Notch1
Log2.FC = -0.58, P-value = 2.76e-03
Hes5
Log2.FC = -0.891, P-value = 2.93e-04
Atoh7
Log2.FC = -0.83, P-value = 1.99e-07
EphB1

**Figure S3. *Pou3f1* is necessary for the induction of the contralateral program**

**(A)** Top: Venn Diagram showing the number of significantly differentially expressed genes (DEGs) in the *Pou3f1* KO E14.5 RNA-Seq that are shared with the cRGC-specific genes identified in a previously published E16.5 cRGC microarray [2]. Bottom: Volcano plot illustrating expression levels of all 38 cRGC-specific genes in the E14.5 *Pou3f1* KO retina. **(B)** Top: Venn Diagram showing the number of DEGs in the *Pou3f1* KO E14.5 RNA-Seq and the significantly down-regulated genes after 48h of Zic2 overexpression [3]. Bottom: Volcano plot illustrating expression levels of all 643 downregulated genes following Zic2 overexpression (OE) at E14.5 in the Pou3f1 KO retina at the same age. **(C)** Venn Diagram showing the number of genes downregulated in Pou3f1 KO at E14.5 and the number of genes upregulated 48h after expression of Pou3f1 at P1. Common 75 genes are considered “Pou3f1-positively regulated genes”. **(D)** Gene expression levels in Log2.FC of the top Pou3f1-positively regulated genes identified in (C). **(E)** Venn Diagram showing the number of genes upregulated in Pou3f1 KO at E14.5 and the number of genes downregulated 48h after expression of Pou3f1 at P1. Common 22 genes are considered “Pou3f1-negatively regulated genes”. **(F)** Gene expression levels in Log2.FC of top Pou3f1-positively regulated genes identified in (E). **(G, H)** Representative gene tracks of upregulated (G) and downregulated genes (H) 48 hours after Pou3f1 overexpression at P1.

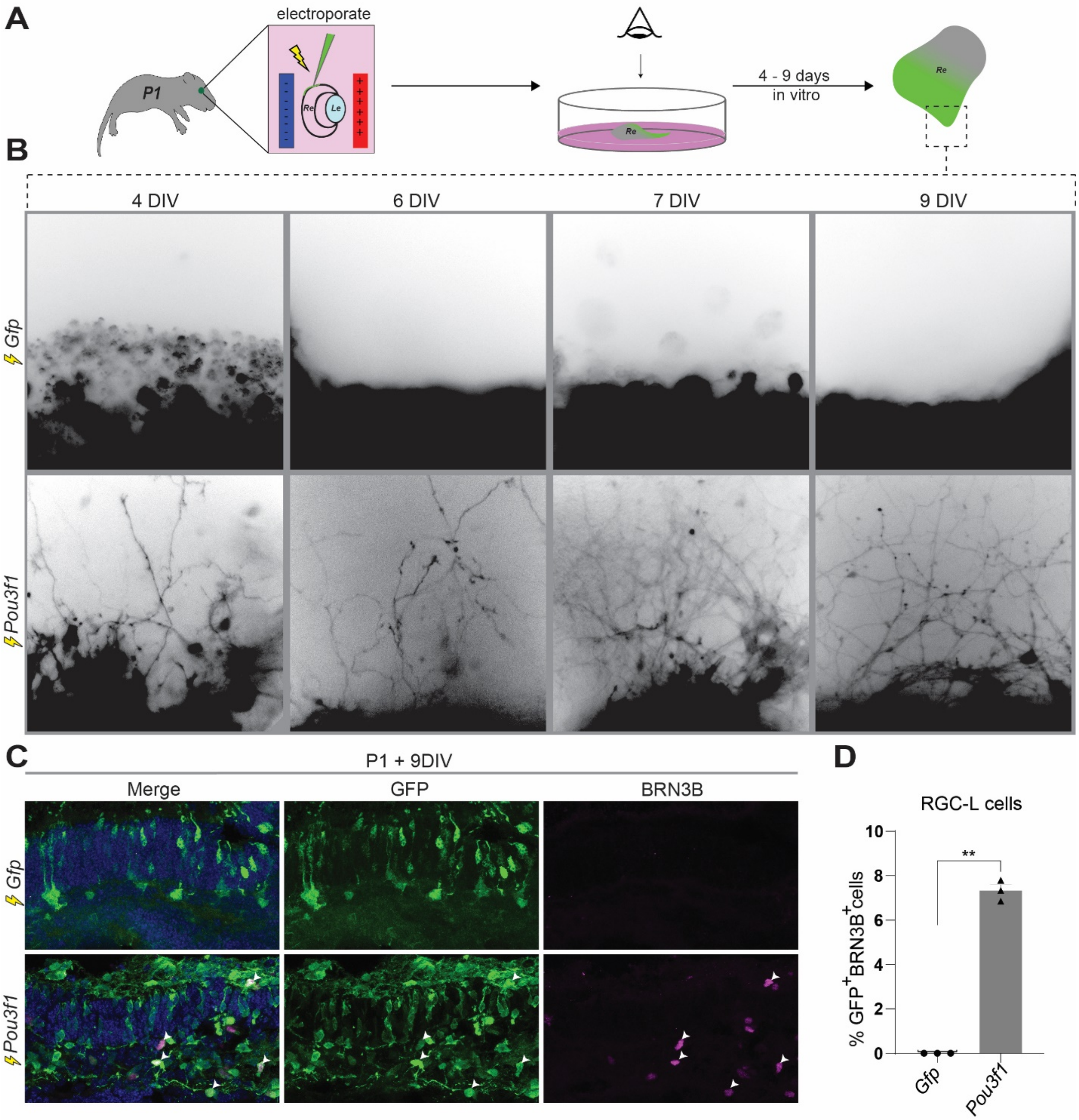
A
electroporate
P1
Re
Le
Re
4 - 9 days
in vitro
Re
B
4 DIV
6 DIV
7 DIV
9 DIV
Gfp
Pou3f1
C
P1 + 9DIV
Merge
GFP
BRN3B
Gfp
Pou3f1
D
RGC-L cells
% GFP+BRN3B+cells
10
8
6
4
2
0
**
Gfp
Pou3f1

**Figure S4.** ***Pou3f1*** **promotes RGC-L cell production** ***ex vivo***

**(A)** Schematic representation of the experimental procedure. Subretinal injections and electroporations were performed on CD1 P1 mouse eyes. Retinas (Re) were separated from the lens (Le) and cultured for up to 9 days. The electroporated patch was imaged on the days listed in (B). **(B)** Imaging of the electroporated patch of GFP+ cells (black) with an inverted fluorescent microscope. Axonal outgrowth in Pou3f1 electroporated explants is observed at 4 DIV and increases in number and complexity by 9 DIV. **(C)** Cross-section and immunostaining for BRN3B of the same electroporated explants shown in (B). Arrowheads highlight $GFP^{+}$/$BRN3B^{+}$ cells. **(D)** Quantification of the proportion of $GFP^{+}$/$Brn3b^{+}$ cells in the *Pou3f1* condition compared to *Gfp* (n = 3).

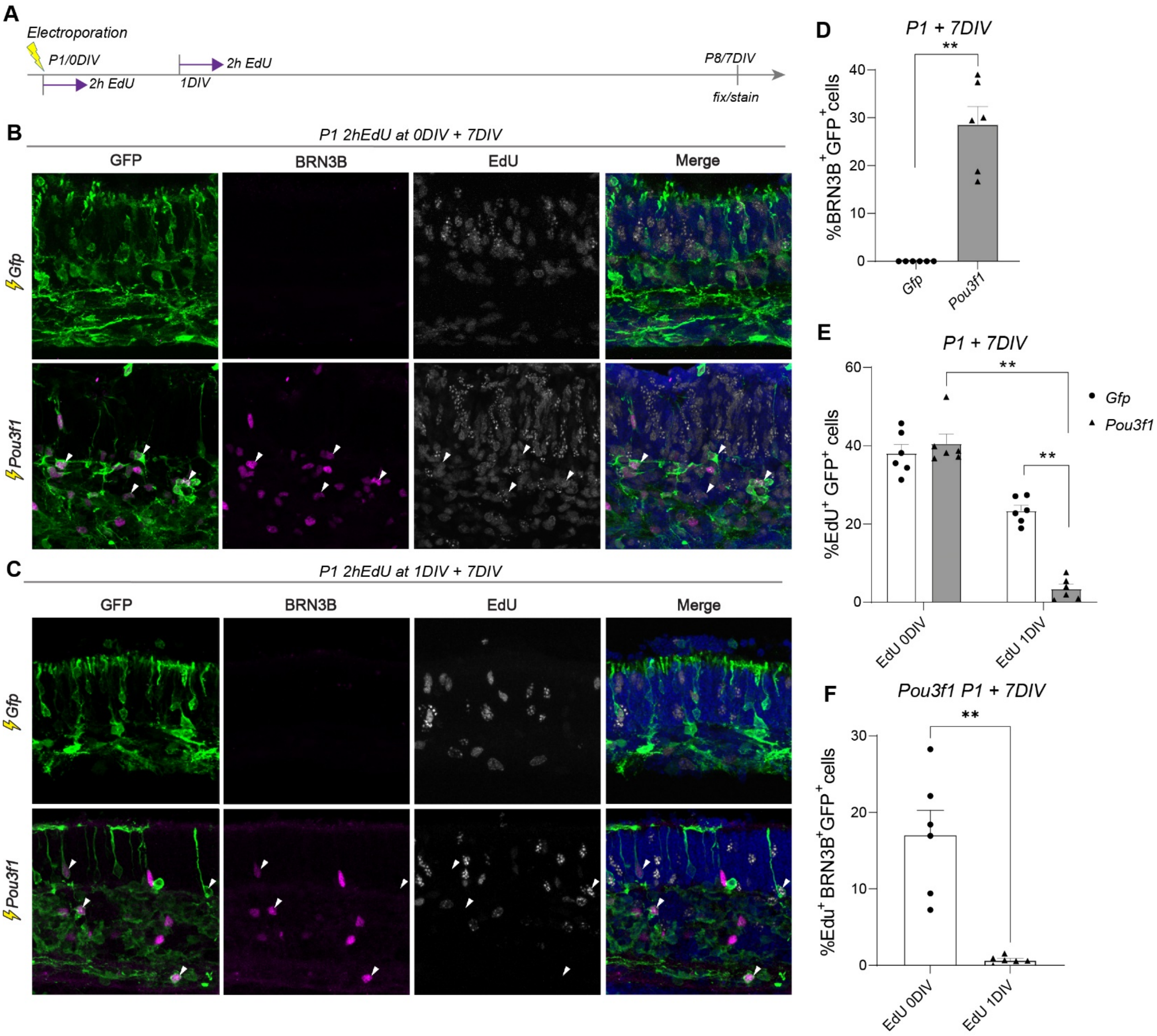


**Figure S5. RGC-L cells originate from retinal progenitor cells**

**(A)** Schematic representation of the experimental procedure. Subretinal electroporation were performed at P1. Retinas were incubated in EdU-containing media for 2h either directly after electroporation (EdU 0DIV) or after 1 day in vitro (EdU, 1DIV). Explants were fixed, sectioned, and stained 7 days after electroporation. **(B, C)** Cross-sections of electroporated explants following EdU pulse at day 0 in vitro (B) or after 1 day in vitro (C). Arrowheads in point to triple positive cells (GFP+/Brn3b+/EdU+) in (B) and GFP+/Brn3b+/EdU- or GFP+/Brn3b-/EdU+ cells in (C). **(D)** Quantification of the proportion of GFP+/Brn3b+ after electroporation of *Gfp* or *Pou3f1* (n = 6, **p<0.01). **(E)** Quantification of the proportion of GFP+/ EdU+ population after *Gfp* or *Pou3f1* electroporation following EdU pulse at 0DIV or 1DIV (n = 6, **p<0.01). **(F)** Quantification of the proportion of GFP+/Brn3b+/EdU+ cell population following EdU pulse at 0DIV and 1DIV in the Pou3f1 electroporated explants (n = 6, **p<0.01).

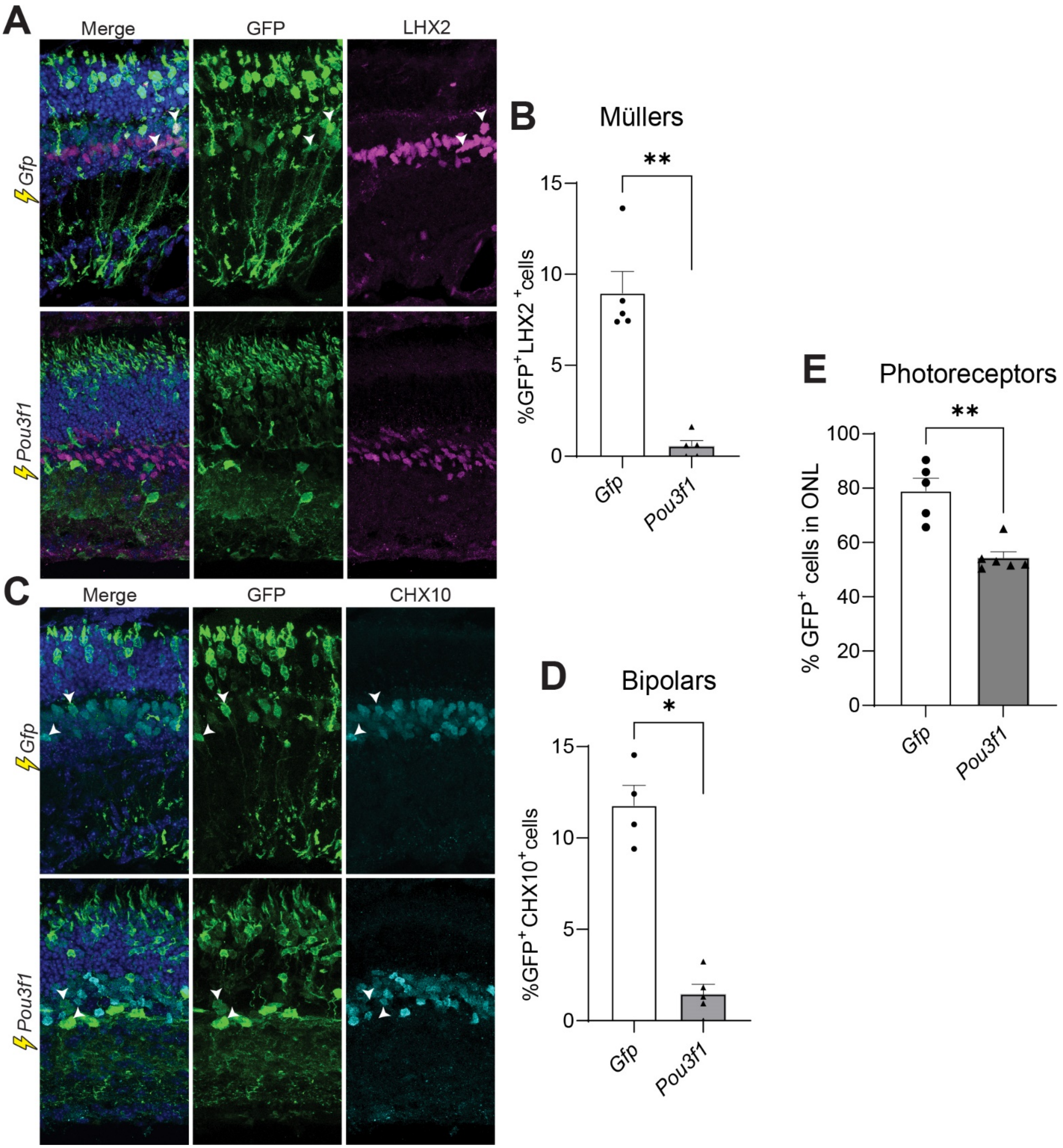

A
Merge
GFP
LHX2
Gfp
Pou3f1
B
Müllers
%GFP+LHX2+cells
**
15
10
5
0
Gfp
Pou3f1
C
Merge
GFP
CHX10
Gfp
Pou3f1
D
Bipolars
%GFP+CHX10+cells
*
15
10
5
0
Gfp
Pou3f1
E
Photoreceptors
% GFP+ cells in ONL
**
100
80
60
40
20
0
Gfp
Pou3f1

**Figure S6. Pou3f1-induced RGC-L cells are produced at the expense of late-born cell types.**

**(A)** Representative images of retinal cross-sections 21 days after electroporation with *Pou3f1* or *Gfp* stained for Lhx2 (Müller cells). **(B)** Quantification of the proportion of GFP+/LHX2+ cells (Müller cells) after *Gfp* or *Pou3f1* electroporation. **(C)** Representative images of retinal cross-sections 21 days after electroporation with *Pou3f1* or *Gfp* stained for Chx10 (bipolar cells) **(D)** Quantification of GFP+/CHX10+ (bipolar cells) after *Gfp*- or *Pou3f1*-electroporation. **(E)** Quantification of the proportion of GFP+ cells in the outer nuclear layer (photoreceptors) after *Gfp* or *Pou3f1* electroporation.

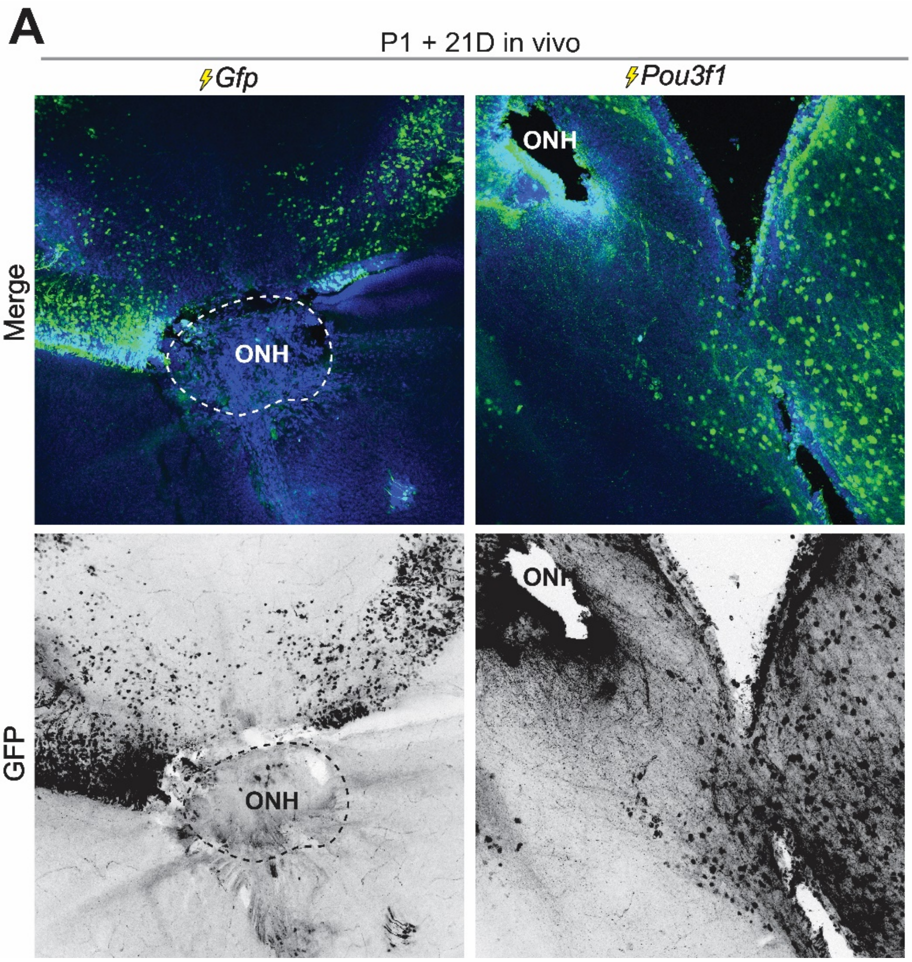


**Figure S7. Pou3f1 expression at P1 in vivo promotes axonal projections towards the optic nerve head**

Representative images of retinal flat mounts 21 days after *Gfp* or *Pou3f1* electroporation at P1. Images in the bottom row are desaturated and inverted to better visualize axonal fasciculation and projection to the optic nerve head (ONH) in the *Pou3f1* condition.

### SUPPLEMENTARY TABLES

**Table S1.** Sequencing reads obtained by RNA-seq 48 hours after electroporation of Pou3f1 in the retina at P1.

**Table S2.** Sequencing reads obtained by RNA-seq 14 days after electroporation of Pou3f1 in the retina at P1.

**Table S3.** Sequencing reads obtained by RNA-seq in *lacZ*+ cells isolated from Pou3f1 HET and Pou3f1 KO.

1. Clark, B.S., et al., *Single-Cell RNA-Seq Analysis of Retinal Development Identifies NFI Factors as Regulating Mitotic Exit and Late-Born Cell Specification.* Neuron, 2019. **102**(6): p. 1111-1126 e5.
2. Wang, Q., et al., *Ipsilateral and Contralateral Retinal Ganglion Cells Express Distinct Genes during Decussation at the Optic Chiasm.* eNeuro, 2016. **3**(6).
3. Escalante, A., et al., *Zic2-dependent axon midline avoidance controls the formation of major ipsilateral tracts in the CNS.* Neuron, 2013. **80**(6): p. 1392-406.
